# A Mediator subunit regulates genome stability via direct and indirect transcriptional targets in *Candida albicans*

**DOI:** 10.64898/2026.11.19.767865

**Authors:** Zinnat Shahina, Philip S. Hendricks, Cristobal Carrera Carriel, Elizabeth Mei, Andrew L. Woodruff, Abigail Ali, Matthew Z. Anderson

## Abstract

Gene families often arise for effector proteins that serve critical functions in organismal survival and fitness. Paralogs encoding the *MED2* subunit of the transcriptional regulatory complex Mediator, designated the telomere-associated (*TLO*) gene family, form the largest lineage-specific gene family expansion in the fungal commensal and opportunistic pathogen *Candida albicans*. Mediator controls global gene expression through site-specific contact with transcription factors in the Tail domain where Tlo resides. The *C. albicans TLO*s have previously been shown to contribute to fungal traits important for growth and host interactions. However, construction of multiple *tlo*Δ/Δ lineages unexpectedly produced strains capable of growth on carbon sources that are toxic to wildtype cells, sorbose and 2-deoxygalactose (2-DOG). Growth on these toxic sugars was partially reversible, suggesting genetic and regulatory contributions to *tlo*Δ/Δ mutant survival. Transcriptional regulation was responsible for growth on sorbose, but survival on 2-DOG resulted from transcriptional regulation and increased genome instability. Survival on 2-DOG was frequently associated with loss of heterozygosity and aneuploidy, and *tlo*Δ/Δ mutants displayed increased telomere dysregulation. Complementation of the *tlo*Δ/Δ mutant with α- and β-*TLO* genes at the *NEUT5L* locus restored susceptibility to both toxic carbon sources and telomere function, whereas addition of most *TLO*γ genes to the *tlo*Δ/Δ mutants did not suppress growth on these media. Taken together, we demonstrate a broad role for these Mediator subunits in controlling cellular functions via direct regulation and genome architecture.

**AUTHORS SUMMARY:** The regulation of gene expression is critical to cell survival when encountering new and diverse environments. In eukaryotes, the Mediator complex is a major source of transcriptional regulation. The fungal organism *Candida albicans* encodes each of the 25 Mediator subunits as a single gene with the exception of a *MED2 homolog* that exists as 14 different copies. These genes are called the te<u>lo</u>mere-associated (*TLO*) gene family, and their collective and individuals roles in the regulation of *C. albicans* gene expression has been difficult to discern. When all of the *TLO* genes were removed from *C. albicans*, the strains were able to grow under conditions that normally kill the fungus. Further investigation found that survival under these toxic conditions was a consequence of direct regulation of genes that promote toxicity and the indirect production of genome-level changes that alter the susceptibility of these cells to killing. These results highlight an unappreciated role for these genes in maintaining the integrity of the *C. albicans* genome in addition to their role in transcriptional regulation.

## Introduction

*Candida albicans* is a polymorphic, commensal yeast that resides on human skin and mucosal surfaces, where it colonizes the oral cavity, gastrointestinal tract, and urogenital tract without causing disease in healthy individuals as part of the normal human microbiota. Each host niche presents distinct environmental conditions that *C. albicans* must respond to. Salivary antimicrobial defences, nutrient fluctuations, and constant exposure to environmental microbes typifies the oral cavity (1); anaerobic, microbiota-rich conditions define the gastrointestinal tract (2); and an acidic pH, sparce microbial landscape, and hormonal fluctuations are common features of the vaginal tract (3). Overgrowth of these niches can lead to life-threatening systemic infection when *C. albicans* access the bloodstream, which is a glucose-rich environment populated by immune cells. Intact host immunity restricts fungal survival in the bloodstream through a combination of oxidative damage, phagocytosis, and nutrient limitation (4).

Successful colonization and pathogenesis by *C. albicans* across host niches requires the remodeling of fungal metabolism, stress tolerance, and morphology between the yeast and filamentous forms (5–7). *C. albicans* overcomes these challenges by adopting distinct transcriptional states and responses to a variety of conditions (8–11). Transcriptional responses are governed by the combined actions of transcription factors and chromatin modifiers to regulate gene expression and do not typically require changes in the genome (12, 13).

The major transcriptional regulatory complex Mediator controls gene expression by bridging transcription factor binding and the recruitment of RNA polymerase. In *C. albicans*, the *MED2* subunit of the Mediator complex underwent a massive lineage-specific expansion from 1 copy in most *Candida* species to 14 copies in the SC5314 genome reference strain (14, 15). These paralogs, named the te<u>lo</u>mere-associated *(TLO)* genes, function as interchangeable Med2 subunits through an N-terminal MED2 domain that anchors them in the Mediator Tail module (16, 17). Sequence variation in the C-terminus of *TLO* genes divides them into three architectural groups (α, β, and γ) that have different levels of expression and subcellular localization (15, 18). Paralogs in each architectural group share ∼97% nucleotide identity, and sequence similarity is lower between groups (roughly 82% identity).

An expanded pool of diverse Tlo proteins has the potential to generate compositionally distinct Mediator complexes. Each Tlo paralogs can be incorporated into a Mediator complex that will interact with distinct transcriptional regulators, control expression of distinct regulons, and generate unique phenotypic profiles (19, 20). Deletion of the complete *C. albicans TLO* gene family resulted in a general fitness defect, including reduced expression of carbohydrate metabolism genes and impaired growth on glucose- and galactose-containing media (19). Restoration of growth on glucose and galactose media in the *tlo*Δ/Δ mutant was dependent on the identity of the reincorporated *TLO* gene. However, this investigation of *TLO* function generally limited analysis to a single representative paralog from each of the three architectural groups. Complementation of *tlo*Δ/Δ strains with different *TLO* genes from the same architectural could produce different phenotypes, suggesting a more detailed analysis of each *TLO* gene would provide greater insight into overlapping and unique paralog functions.

The previously constructed *tlo*Δ/Δ mutants also displayed reduced survival under multiple stress conditions with the exception of azole antifungal drugs (19, 21). Loss of the *TLO* genes altered the sterol composition of the fungal cell membrane and increased azole drug tolerance. Resistance to azole antifungal drugs by *C. albicans* is commonly conferred by karyotypic changes, including trisomy for multiple different chromosomes that encode the drug target or regulators of drug efflux pump expression (22, 23). Genes encoded on trisomic chromosomes increase in expression in accordance with their dosage, thereby promoting resistance (22, 24–26). *C. albicans* chromosomes can also undergo whole or partial loss of heterozygosity (LOH), in which alleles from one chromosome are replaced by alleles from the other homolog (27, 28). The simultaneous fixation of many alleles across a chromosome and reduction in standing genetic diversity can have profound impacts on cell physiology and behaviors (29, 30).

Early investigations into growth on toxic carbon sources highlighted the role of aneuploidy and LOH towards cell survival. Growth on L-sorbose interferes with *C. albicans* cell wall synthesis and leads to cell death (31, 32). However, monosomy of chromosome 5 (Chr5) relieves repression of the *SOU1* gene on Chr4 and increases conversion of L-sorbose to D-sorbitol (33), neutralizing its toxicity. Increased copies of Chr4 likewise promote L-sorbose survival by increasing *SOU1* copy number and gene expression directly (34–36). In contrast to L-sorbose, a metabolic intermediate of 2-deoxygalactose (2-DOG), 2-deoxy-galactose-1-phosphate, and not the sugar itself produces toxicity by blocking glycolysis (37).Loss of the *GAL1* galactokinase either through LOH at a hemizygous locus or construction of a targeted deletion confers 2-DOG resistance (38–40).

Genes that would promote resistance to toxic compounds or antifungal drugs through chromosome instability (CIN) typically function in DNA repair and cell cycle progression (41–46). Mutants in genes that promote CIN rarely produce specific phenotypes. Interference with coordination between nuclear segregation and cell division will produce a range of diverse karyotypes that are not limited to a single chromosome or ploidy state. Although transcriptional regulators are likely involved in this coordination, substantially less is known about their involvement in CIN. Mutants for *S. cerevisiae MED9*, a subunit of the Middle module of Mediator, increase DNA segregation defects (*47*), but no other roles for Mediator in CIN have been described. Therefore, gaps exist in our knowledge to connect transcriptional regulation to CIN and the resulting karyotypic variation that promote cell survival.

In this study, we set out to examine the contribution of individual *TLO* genes as transcriptional regulators to genome variation. We begun this work by investigating the ability for *tlo*Δ/Δ mutants to survive in the presence of the toxic carbon sources, L-sorbose and 2-DOG. In contrast to wildtype cells, *tlo*Δ/Δ mutants survived on these sugars at high frequencies. Survival on sorbose-containing medium functioned primarily through transcriptional control of target genes. In contrast, growth on 2-DOG resulted from a combination of transcriptional regulation by *TLO* genes and increased genome instability via aneuploidy and LOH. The wildtype phenotypes were largely restored by complementation of the *tlo*Δ/Δ mutant with *TLO*α and *TLO*β genes but not *TLO*γ paralogs, with the exception of *TLO*γ*4*. Taken together, these results reveal a role for *TLO* genes in promoting cell survival through a combination of direct transcriptional regulation of target genes and indirect control of broader genome integrity that is largely compartmentalized along architectural groups.

## MATERIALS AND METHODS

### Media and reagents

Yeast extract-peptone-dextrose (YPD - 1% yeast extract, 2% peptone, and 2% dextrose) and synthetic complete defined (SCD – 0.7% Yeast Nitrogen Base (YNB) w/o amino acids, w/o carbohydrates, 0.17% Complete Amino Acid Drop Out and 2% dextrose) media were prepared by dissolving the appropriate nutrient into Type 1 laboratory-grade water and autoclaving as previously described (48).Yeast extract–peptone–maltose (YPM) medium was prepared as described above for YPD with maltose replacing dextrose as the carbon source. YPD and SCD containing 200 µg ml^−1^ nourseothricin (Werner Bioagents, Jena, Germany) were used to select for nourseothricin-resistant (NAT^R^) strains. SCD medium lacking L-leucine (SCD-LEU) was used to select for leucine prototrophic strains. 2-deoxygalactose (2-DOG) medium consisted of synthetic complete (SC) medium supplemented with 2% glycerol, 2 mg/mL 2-DOG (99%, Thermofisher, Waltham, MA, USA), and 2% agar. L-sorbose and glucose plates were made with 0.7% yeast nitrogen base without amino acid, autoclaved, and combined with filter sterilized 2% L-Sorbose (MilliporeSigma, Burlington, USA) or 2% Glucose and 10 mg/ml uridine, respectively, before pouring into petri dishes.

### Plasmid construction

Plasmids containing single *TLO* genes were constructed using a modified gap repair cloning protocol (49) and are listed in Table S1. Briefly, two sets of 40 bp reverse complement primers were designed. One set amplified the *TLO* gene from the genome of the parental *LEU2*/*leu2* strain from at least 400 bp upstream of the start codon to the codon prior to the stop codon with 20 bp overhangs corresponding to the second primer set. The second primer set amplified a plasmid vector with a 6xHis-1xFLAG *SAT1* FLPer cassette with overhangs matching the amplified *TLO* gene. DNA targets were amplified using Repliqa HiFi ToughMix (Quantabio) and amplification products were purified using the SV Wizard PCR cleanup kit (Promega). The purified products were transformed into DH5α *E. coli* (Thermofisher,Waltham, MA, USA) in a 2:1 molar insert to vector fragment ratio with 150 ng of vector used, and successful repair products were selected for on LB+ 50 µg/mL chloramphenicol. Gene sequence and vector insertion were confirmed by whole plasmid sequencing by the University of Wisconsin Biotech Center (UWBC). This method was used to construct all plasmids used in this study except the *TLO2* and *TLO10* containing-plasmids.

### *De novo* plasmid synthesis

The desired plasmids for *TLO*β*2* and *TLO*α*10* could not be obtained from gap repair cloning, so these plasmids were synthesized by the Thermofisher GeneArt gene synthesis service. The full *TLO* coding sequence and upstream intergenic sequence was included in the same 6xHis-1xFLAG *SAT1* FLP plasmid as the other *TLO* genes. The *TLO*β*2* plasmid contained an ampicillin resistance cassette, and the *TLO*α*10* plasmid contained a kanamycin resistance cassette.

### Strain construction

All strains are described in Table S2. A diploid strain derived from the Chr5 trisomic AHY940, which is a heterozygous knockout for the *LEU2* gene, was used as the parental strain for two independent CRISPR-Cas9 deletions of *TLOα34* using the deletion method described in (50). Primers were designed to generate a guide RNA (gRNA) that was specific to the region immediately upstream of the *TLOα34* coding sequence (CDS) and a donor DNA (dDNA) was designed to facilitate removal of *TLOα34*. Successful deletions were confirmed first by PCR and then by whole genome sequencing (WGS) performed at the Ohio State University Applied Microbiology Sequencing Laboratory (AMSL). These strains were then used as the platform for CRISPR-Cas9 deletion of the remaining *TLO* genes. Deletion of the remaining 13 *TLO* genes was performed in a single step using CRISPR-Cas9 as described above. Two gRNAs targeted a conserved region of the coding sequences, and eight times the recommended amount of dDNA was included in the transformation to increase the availability of the repair template. The dDNA was designed to remove the full coding sequence of the *TLOγ* genes and all but the last 120 base pairs (bp) of the *TLOα* and *TLOβ* genes. Successful deletion of all *TLO*s was confirmed using universal *TLO* primers and then by WGS provided by AMSL. Two independent *tlo*α*34*Δ/Δ mutants were used to generate two independent complete *TLO* knockout (*tlo*Δ/Δ) strains. All strains were grown on SCD-leucine media post transformation to remove the CRISPR-Cas9 system from the genome.

Other homozygous deletion mutants were produced using the LEUpOUT method of CRISPR-Cas9 designed for *C. albicans* (50). All guide RNAs and primers used for gene deletions are listed in Table S3.

To generate *SOU1* heterozygous strain, an amplicon was generated from pSFS2a (51) containing the NAT-resistance locus *SAT1* and directed to the *SOU1* locus by 70bp long oligo overhangs using a standard lithium acetate transformation (52). After verification of *SOU1* heterozygosity via PCR, the *SAT1*-FLP cassette was recycled by plating to YPM+20 µg/mL NAT as described above.

Reintegration *of TLO* genes into the *tlo*Δ/Δ mutants was performed via long oligo mediated lithium acetate transformation. A set of 90 bp primers were designed to amplify the tagged *TLO* gene and removable drug resistance cassette from the plasmid. The primers had 70 bp of homology to the *NEUT5L* locus to target constructs to this neutral locus. Plasmids were amplified with Repliqa Hifi ToughMix, and four separate reactions were combined and purified with SeraMag magnetic beads. The *tlo*Δ/Δ mutants were grown overnight to an OD between 4-8 in YPD at 25°C. Approximately 2.0×10^8 cells were centrifuged and washed once with water. Cells were then suspended in 240 µL of TELiOAc master mix (100 μL 10x Tris-EDTA pH=8, 100 μL 1M Lithium Acetate, 20 μL 1M DTT, and 20 μL of 20 mg/mL boiled sheared salmon sperm), 800 µL of 50% PEG 3350, and 3 µg of the amplified insertion cassette and mixed via pipetting.

The transformation reactions were then incubated for 1 hour at 30*C and then 45 minutes at 42*C. The reactions were then briefly centrifuged, and the supernatant was discarded. The pellet was resuspended in 2 mL YPD and incubated at 30*C for 4 hours. Cells were plated to YPD+20 µg/mL nourseothricin (NAT) to select for resistance colonies. Successful transformants were identified by PCR, and the resistance cassette was removed by plating on YPM+20 µg/mL NAT. Small colonies had lost the resistance cassette and were confirmed by patching cells to YPD and YPD+NAT. Reintegrant strains for each *TLO* were constructed in both *tlo*Δ/Δ lineages for two independent strains. Strains were whole genome sequenced using 2×150 paired Illumina short read sequencing provided by the UWBC and checked using YMAP (53) to confirm no large LOH events or aneuploidies occurred during strain construction.

### Spot assay

Cells were struck onto YPD plates from glycerol stocks at −80°C and grown at 30°C for 2 days. Cells were collected from the plate and washed thrice with PBS to remove traces of glucose. Cells were counted using a hemocytometer, resuspended at a concentration of approximately 10^6^ - 10^8^ cells/mL, serially diluted in 10-fold increments, and 6 µL of each dilution was spotted onto the corresponding plates. Spot assays used 2-DOG, L-sorbose, SC+glucose, and YPD plates. Spotted plates were incubated at 30 °C for 2 days (SC+glucose and YPD) or 4-5 days (2-DOG and L-sorbose)and imaged using a ChemiDoc XRS+ molecular imager (Bio-Rad, Hercules, CA, USA) (27, 54, 55).

### Fluctuation analysis

Cells from each strain were taken from YPD plates, washed with PBS 3x, counted using a hemocytometer, and 100 colonies were plated to YPD and 2-DOG. Plates were incubated at 30°C for 2–3 days and the number of colonies were counted on each medium. Images were taken using a ChemiDoc XRS+ molecular imager (Bio-Rad, Hercules, CA, USA).

To distinguish loss of the *GAL1* gene from silencing, single colonies from 2-DOG plates were replica plated on YPD and incubated for 24 h at 30°C. YPD plates were replica plated back to 2-DOG medium and incubated for 4 days at 30°C. Colonies were enumerated on 2-DOG plates to determine the frequency of *GAL1* loss and silencing.

### Quantitative reverse transcription PCR (qRT-PCR)

Expression of genes of interest was measured by collecting total RNA from colonies grown on solid medium, YPD, galactose after 2 days and 2-DOG after 6 days at 30°RNA was harvested using the MasterPure Yeast RNA Purification kit (Biosearch Technologies, Middleton, WI, USA) according to the manufacturer’s instructions and subsequently treated with DNase I. RNA concentrations were measured using a NanoDrop OneC Microvolume UV-Vis Spectrophotometer (Fisher Scientific, Waltham, MA, USA). cDNA was generated from 1 µg of DNase I treated RNA using oligo(dT)18 and SuperScript III Reverse Transcriptase (Thermo Scientific, Waltham, MA, USA). The cDNA was screened for the presence of genomic DNA contamination via PCR using an intron-spanning primer set for ribosomal protein large subunit 6 (RPL6) listed in Table S3 (15, 56).

The qRT-PCR reaction was performed with 50 ng of cDNA in a 12 µl reaction with PowerUp SYBR Green (Applied Biosystems, Foster City, CA, USA) using an Applied Biosystems QuantStudio 5 Real-Time PCR System. Oligos used for qRT-PCR are provided in Table S3. Amplification was performed for 40 cycles (thermal cycling conditions: Initial steps of 50°C, 2 minutes and 95 °C, 2 minutes; and then, 40 cycles of 95 °C, 15 sec; 60 °C, 15 sec and 72 °C for 1 minute). To verify the specificity and purity of the amplified DNA product, melt curves were assessed. The Ct values for each experiment were recorded, and gene expression was calculated using the 2−ΔCt method, where expression of each gene was normalized to *ACT1* expression. Experiments for each gene were performed with a minimum of four biological replicates per genotype with two technical replicates each. Each biological replicate was an independent colony or patch. Primers were validated using standard curves.

### Quantitative PCR

Quantitative PCR was performed as described above for qRT-PCR from gDNA isolated using the Zymogen Quick-DNA Fungal/Bacterial Miniprep kit (Zymogen). 50 ng of DNA was used per reaction with the qPCR primers listed in Table S3. The relative abundance of the target gene was determined in comparison to *ACT1*. Production of single amplification products was verified using melt curve analysis.

Oligos matching the 23-bp *Candida* telomeric repeat was used to measure telomere abundance (57). Quantitative PCR was performed in 20 µL reactions using 10 µL of SYBR Green (Applied Biosystems, Foster City, CA, USA),1 µL of primer 1 (0.5 µM),1 µL of primer 2 (0.5 µM), 50 ng of DNA, and dH_2_O using an Applied Biosystems QuantStudio 5 Real-Time PCR System with the oligos provided in Table S3 for 40 cycles (thermal cycling conditions: Initial steps of 95 °C, 10 minutes; and then, 40 cycles of 95 °C,15 sec; 55 °C, 30 sec and 72 °C for 30 sec). Melt curve was used to verify the specificity and purity of the amplified DNA product.

### Growth curve assays

Colonies were picked from YPD plates and grown overnight at 30 °C in 300 µL of liquid YPD medium. Overnight cultures were arrayed in a 96 deep-well plate with shaking at 125 revolutions per minute (rpm). After growing overnight, saturated cultures were diluted 1 : 40 into H_2_O and then 1 : 50 into 150 uL of fresh liquid medium in a flat-bottom 96-well plate. Media types for growth curves included YPD, minimal medium + 2% L-sorbose, or minimal medium + 2% glucose. Plates were sealed with a sterile, optically transparent polyester adhesive sealing film and incubated at 30°C with continuous shaking using a Synergy H1 microplate reader (Biotek, Shoreline, WA). The OD_600_ was recorded every 15 minutes for 48 - 144 hours. Data were analysed using GrowthCurveR (58) and graphs generated on GraphPad Prism (v6). All assays included four biological replicates, each with two technical replicates.

### *In vitro* evolution and whole genome sequencing

*Candida* cells were passaged on solid rich medium (YPD) agar plates at 30°C. Strains were re-struck to fresh plates every 2 days for 20 days from the thick part of the streak, which minimizes bottlenecking of the population (59).Single colonies were obtained from the final streaks, and genomic DNA was collected from each colony. Zymogen Quick-DNA Fungal/Bacterial Miniprep kit (Zymogen) was used for DNA extraction, and DNA concentrations were quantified using the Qubit dsDNA Broad Range Assay kit (Thermo Fisher, Waltham, MA, USA). DNA was sent to University of Wisconsin-Madison Biotechnology Center for library preparation and sequencing. All colony libraries were sequenced as 150 basepair paired-end reads on an Illumina NovaSeq to an average depth of 30 million.

### Genomic variants analysis

The quality of the raw reads was assessed with FastQC (v0.12.1), and trimmed with fastp (0.24.0) to remove low-quality bases and adapters using the pair-end adapter detection option. We then removed the first 20 bases from the 5’ end and 5 bases from the 3’ end to further remove low-quality regions.

Trimmed reads were mapped against the SC5314 reference genome (version A21-s02-m09-r12) using Bowtie 2 (v2.2.6–2) (60). Aligned SAM files were then converted to BAM format, sorted, and indexed using samtools v1.7 (61). Read group information was added using the Genome Analysis Toolkit (GATK v3.167, Picard) AddOrReplaceReadGroups module (62), and potential PCR duplicates were marked using MarkDuplicate. Variant calling was performed with GATk HaplotypeCaller using genomic variant call format (GVCF) mode. Individual GVCF files were combined using CombineGVCFs, followed by genotyping with GenotypeGVCFs. We then filtered low-quality variants using VariantFiltration with the following thresholds: quality by depth (QD < 2.0), variant quality (QUAL < 50), Fisher strand bias (FS > 60.0), mapping quality (MQ < 40.0), mapping quality rank sum (MQRankSum < –12.5), read position rank sum (ReadPosRankSum < −8), and site-level read depth (DP < 20). Variants that did not pass the filtering thresholds were excluded using SelectVariants. The module GATK VariantsToTable was used to export the remaining variants to a tab-delimited table.

Colony-specific variants were identified using a custom Python script (Python 3.13.2, Pandas 2.3.2). For each sample, variants were assigned a unique ID based on chromosome, genomic coordinate, reference allele, and alternate allele. To exclude preexisting genetic differences, variants also seen in the colony’s corresponding background strain were filtered out. The remaining variants were then compared among colonies of the same strain to identify colony-specific variants. A variant was considered colony-specific if an alternate allele at a genomic position was detected in only one colony within that strain. Variants shared between colonies were excluded from the analysis. Variants located in the mitochondrial genome were also excluded.

### GREP analysis of telomeric repeats

Reads were isolated from the FASTQ file for each passaged strain that contained the telomeric repeat sequence“GTACACCAAGAAGTTAGACATCC|GGATGTCTAACTTCTTGGTGTAC” after filtering for read quality (63). Differences in telomere read abundance were calculated as a ratio to the total read count for a sample and plotted in GraphPad Prism.

### Chronological lifespan (CLS) analyses

The CLS assay was carried out as previously described with slight modification (64–66). Briefly, overnight cultures were grown in SCD medium to saturation and diluted the next day into 10 mLs of fresh SCD medium to an OD_600_ of ∼0.1 (1:100 dilution). Cultures were incubated at 30°C with shaking (220 rpm) throughout the rest of the experiment where day 0 marks the point of initial inoculation. Cells were assessed for viability for days 2 to 14 and day 17 used propidium iodide (PI) staining.

To stain cells with PI, aliquots were taken from the shaking culture, were washed with 3x PBS, PI was added to a final concentration of 1 µg/mL. Cells were then incubated for 30 minutes at room temperature in the dark before being imaged by epifluorescence (λ_ex_ = 493 nm; λ_em_ = 636 nm) at 63× magnification. Five fields of view were used for analysis per biological replicate and included at least 500 cells. Viability was plotted over time with day 0 being set as 100% viability.

### Chromatin immunoprecipitation followed by sequencing

Three independent cultures of each Tlo-6xHIS/FLAG (experimental) and Nup49-6xHIS/FLAG (control) strain were grown overnight in 2 ml YPD at 30°C (Table S2). Saturated cultures were diluted to an OD_600_ of 0.2 in 50 mLs of YPD and grown at 30°C to an OD_600_ of 1. DNA cross-linking, DNA shearing and chromatin immunoprecipitation (ChIP) were conducted as described in (67) with some modifications. Briefly, cultures were spun at 3800 rpm for 3 minutes, washed with 5ml of PBS, re-suspended in 36 mLs PBS, and fixed with 1 ml of 37% formaldehyde (final concentration, 1%). Cells were incubated at 30°C for 30 minutes with agitation with formaldehyde before being quenched by adding 2 mLs of 2.5 M glycine and incubating for 10 minutes. Cells were spun down, washed with 1 ml of ice-cold PBS, and snap-frozen in liquid nitrogen. Total cell extracts were prepared by adding 400 uL of FA lysis buffer (50 mM HEPES pH 7.5, 140 mM NaCl, 1 mM EDTA, 1% Triton X-100, 0.1% Na deoxycholate,), 4 uL of protease inhibitor (Pierce™ Protease Inhibitor Mini Tablets, EDTA-free), 2 mM (3.2 µL) of phenylmethylsulfonyl fluoride (PMSF, Thermofisher, Waltham, MA, USA), and an equal volume (∼500 uL) of acid-washed glass beads (MilliporeSigma, Burlington, USA). Tubes underwent bead beating for 7 cycles (setting 4 for 30s per cycle with 1 minute on ice between cycle) using a Fisherbrand™ Bead Mill 24 Homogenizer (Waltham, MA, USA). Extracts were then sonicated with 14 rounds of 10s pulse at 40 microns and 1 minute on ice (Fisherbrand model 50 sonic dismembrator with 1/8th inch tip, Waltham, MA, USA). DNA was assessed by gel electrophoresis for fragment lengths. This approach yielded 300-500-bp fragments on average.

Immunoprecipitation was performed by incubating the sonicated extracts overnight with Monoclonal ANTI-FLAG M2 antibody at 4°C with rocking. The samples were then coupled to Pierce Protein A/G Magnetic Beads (Thermo Scientific, Waltham, MA, USA) at 4°C and 16 rpm for 2 hours. Antibody-bead complexes were washed once with FA lysis buffer without PI, twice with wash buffer-1 supplemented with 0.5M NaCl, twice with wash buffer-2 (10 mM Tris-HCl, pH 7.5, 0.2M LiCl, 1 mM EDTA, 0.5% Na deoxycholate), and once with TE buffer (10 mM Tris, pH 7.5, 1 mM EDTA). DNA was eluted from the beads with ChIP elution buffer (SantaCruzBiotech, Dallas, TX, USA) by first incubating at 65°C for 2 hours, followed by rotation at 16 rpm for 10 minutes at RT, and centrifugation at 13,000 rpm for 2 minutes. The eluted material was then treated with pronase in a thermocycler using the following conditions (42°C – 2h, 65°C – 6h, Hold at 10°C). DNA was purified by magnetic beads and resuspended in 25 μL of TE. Final DNA concentrations ranged from 0.6-0.9 ng/µL. Library construction was performed with 1 ng of the immunoprecipitated DNA with the NEBNext Ultra II DNA Library Prep Kit as recommended by the manufacturer (NEB, Ipswich, MA, USA). Libraries were verified using an Agilent 2100 bioanalyzer (Agilent Technologies, Santa Clara, CA, USA). Prepared libraries were then sent to University of Wisconsin-Madison Biotechnology center and were sequenced as 150 bp paired-end reads on an Illumina NovaSeq to a depth of at least 37.5 million reads/sample.

Peak enrichment was performed using the model-based analysis of ChIP-seq (MACS3) (68). Samples were first assessed by FastQC (v0.12.1), adaptor sequences and orphan reads were removed using FASTP (v0.24.0) (69), and reassessed with FastQC. Reads were aligned to the SC5314 reference genome (version A21-s02-m09-r12) obtained from the *Candida* Genome Database (http://www.candidagenome.org) using Bowtie2 (2.5.2) (60). Aligned reads were converted to the BAM format, sorted, and indexed using SAMtools (v1.9) (61). Conversion to bigwig files was performed using deepTools bamCoverage (v3.5.6) with a bin size of 10 and normalization to genomic context. Peak calling was performed with MACS3 callpeak (v3.0.3) using standard parameters for broad regions and normalized to the effective genome size with the Nup49-6xHIS/FLAG also serving as a non-specific binding control. Integrated genome viewer (IGV) (v2.18.2) was used to visualize and evaluate peaks and alignments.

### Data availability

Sequencing data generated as part of this analysis is available in the Sequence Read Archive (SRA) under PRJNA1543763.

### Statistical analysis

Statistical analyses were conducted using GraphPad Prism (Version 6.0; La Jolla, CA, USA). Unless otherwise stated, the error bars indicate the standard deviation of the mean among biological replicates.

## Results

### Deletion of the *C. albicans TLO* genes allows for growth on toxic carbon sources

Previously, we constructed two *tlo*Δ/Δ lineages using CRISPR-Cas9 by targeting the conserved MED2 domain of this expanded gene family (70).The *tlo*Δ/Δ mutant colonies were ruffled on YPD medium at 30°C and contained a mixture of yeast, pseudohyphae, and hyphae in contrast to the smooth colonies of the wildtype parent that only contained yeast cells (Figure 1a). The altered cell and colony morphology of the *tlo*Δ/Δ mutant was consistent with the phenotype of a previously obtained *tlo*Δ/Δ mutant that contained multiple chromosomal rearrangements (19). Thus, the altered cellular and colony morphology of the *tlo*Δ/Δ mutants is likely a consequence of gene family loss and not karyotypic rearrangements following the use of CRISPR-Cas9.

**Figure 1.**
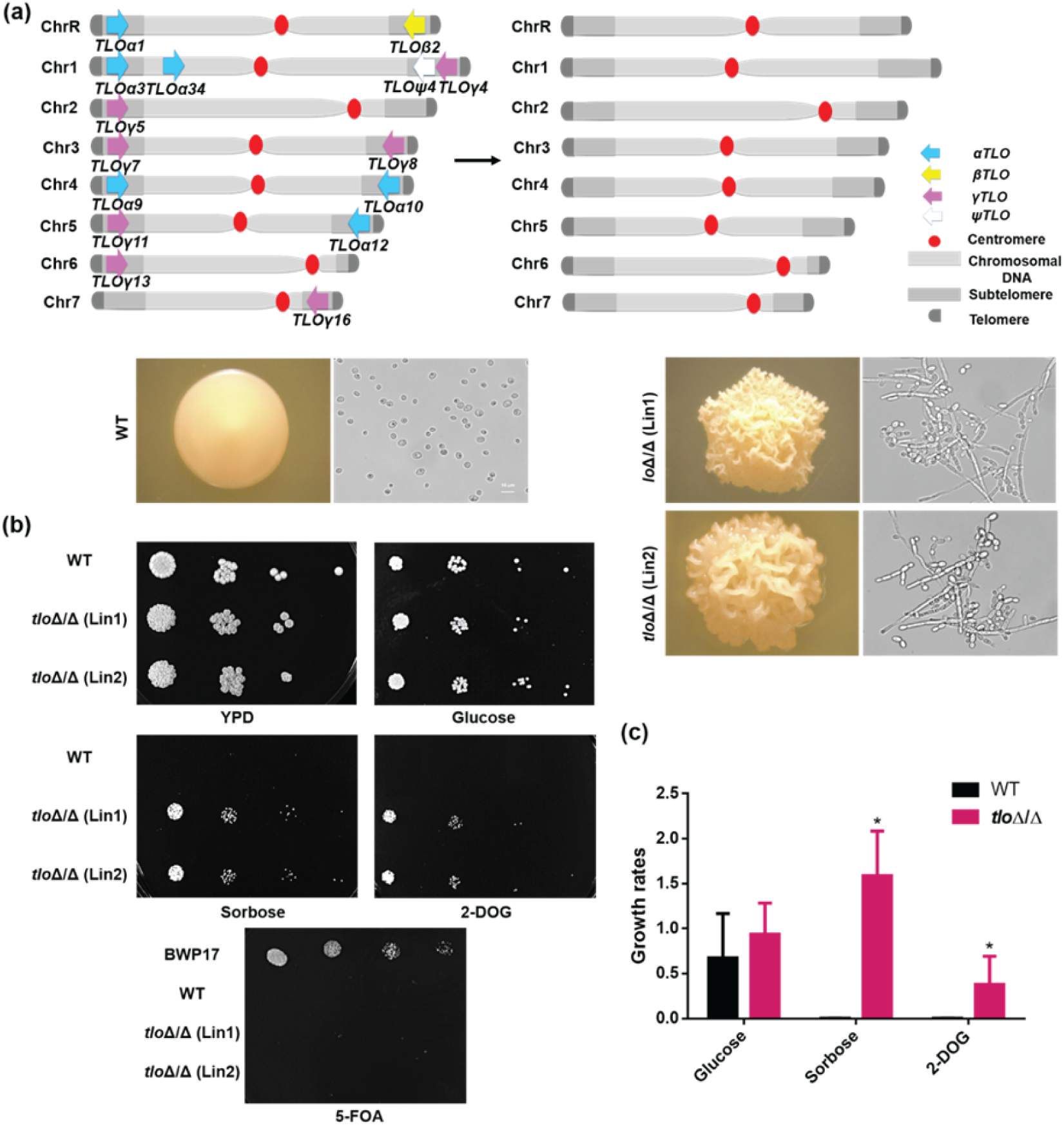
Loss of *TLO*s facilitates growth on toxic carbon sources. (**a**) CRISPR-Cas9 was used to construct *tlo* null mutants in *C. albicans* strain SC5314. Loss of *TLO* genes produced cell populations with wrinkled colonies dominated by pseudohyphae in contrast to the smooth colonies dominated by yeast cells in the wildtype (WT) strain. (**b**) WT and *tlo*Δ/Δ mutants were spotted in 10-fold dilutions onto YPD and synthetic complete medium with 2% glucose, 2% L-sorbose, or 0.2% 2-DOG. Colonies were allowed to grow for 6 days at 30°C before imaging. (**c**) Quantification of spot dilution colony counts. N=3. Statistical significance was determined by Kruskal-Wallis and a Dunnett’s post hoc test v. WT. * p<0.05

Although Mediator has a broad role in gene regulation, little evidence links it to chromosome instability. To test for genome instability phenotypes linked to *C. albicans TLO* genes, cells from the wildtype parental strains and two *tlo*Δ/Δ mutants were plated to toxic substrates, including sorbose, 2-deoxygalactose (2-DOG), and 5-fluoroorotic acid (5-FOA). 5-FOA is a chemical analog of uracil that produces a toxic intermediate compound, 5-fluorouracil, when *de novo* uracil biosynthesis is intact. Inactivation of *URA3* disrupts uracil biosynthesis and provides resistance to 5-FOA (71). An SC5314-derived strain containing a homozygous deletion of *URA3* grew on 5-FOA plates as expected, but neither the SC5314 wildtype nor the *tlo*Δ/Δ mutants formed colonies (Figure 1b). Surprisingly, the *tlo*Δ/Δ mutants grew robustly on sorbose and 2-DOG media, whereas no growth was observed for wildtype cells (Figure 1b,c). This suggests that loss of *TLO* genes affected specific loci or pathways and does not promote growth of toxic carbon sources broadly.

### *TLO*s control sorbose survival through regulation of *CSU* genes

To investigate the mechanism promoting survival of the *tlo*Δ/Δ mutants on sorbose, wildtype and *tlo*Δ/Δ cells were plated to rich media (YPD and YNB+2% glucose) and YNB+2% sorbose. As previously observed, all strains grew on rich media, but only the *tlo*Δ/Δ mutants could grow on sorbose medium (Figure 2a,b). The improved growth of the *tlo*Δ/Δ mutants in sorbose was not due to a general increase in growth rates. Instead, the *tlo*Δ/Δ mutants displayed a roughly 2-fold slower doubling time in liquid YPD and YNB+2% glucose at 30°C compared to the wildtype strain (Figure 2c).

**Figure 2.**
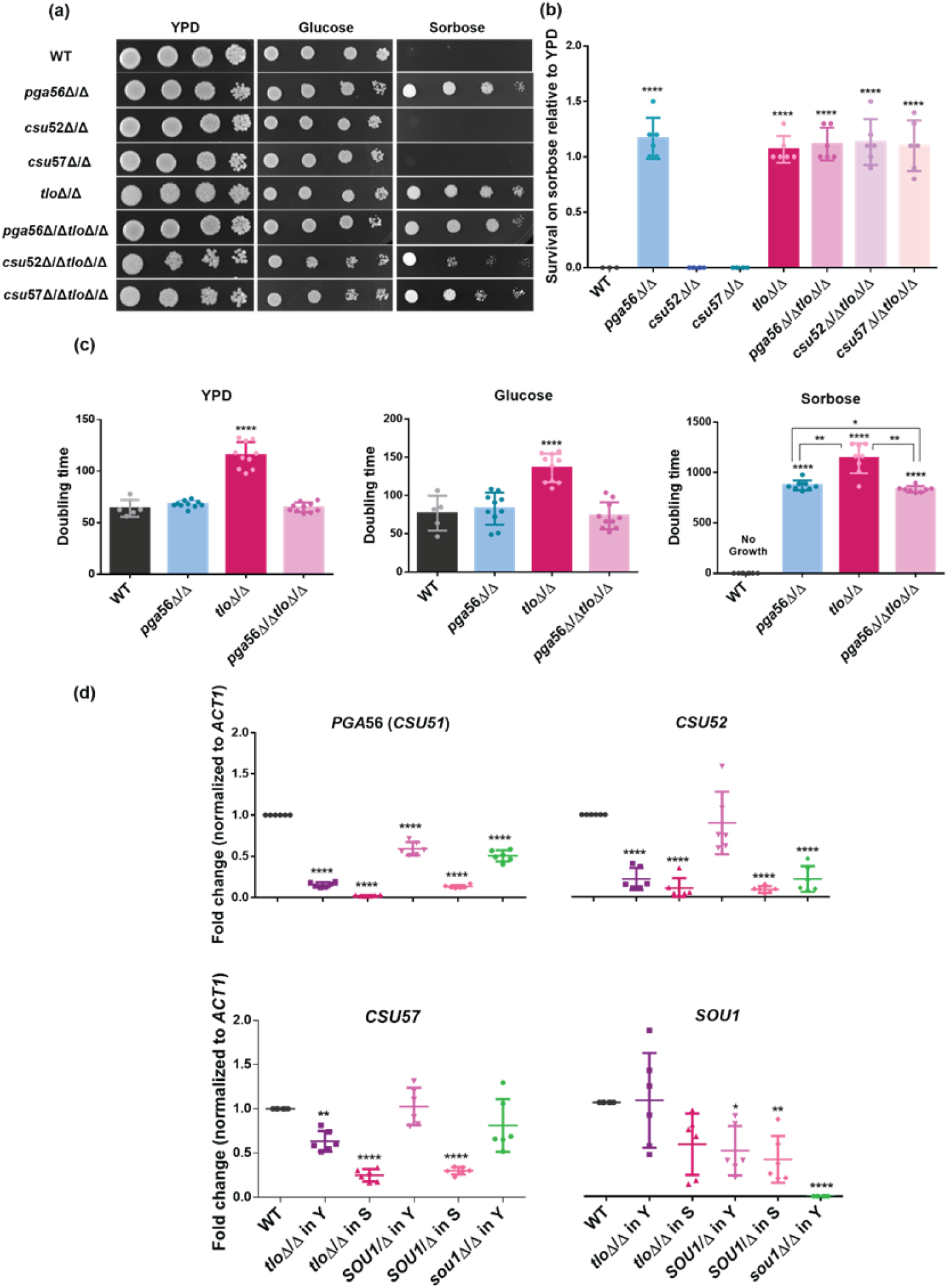
Growth of *tlo*Δ/Δ cells on sorbose is associated with decreased expression of sorbose utilization genes. (**a**) Cells of the indicated genotypes were spotted in 10-fold dilutions starting with 10^8^ cells on the left onto YPD and yeast nitrogen base (YNB) medium supplemented with 2% glucose or L-sorbose. Colonies were allowed to grow for 6 days at 30°C before imaging. (**b**) One hundred cells from each strain were plated onto YNB+2% L-sorbose or YPD, allowed to grow for six days, and the resulting colonies were quantified. N=3. (**c**) Transcript abundance of sorbose utilization genes was measured by qRT-PCR in the wild-type and *tlo*Δ/Δ mutants and normalized to *ACT1*. Y and S indicate growth on solid YPD or sorbose agar medium at 30°C for 2 or 6 days, respectively. N=6. (**d**) Wild-type and the indicated mutant strains were grown in liquid YPD, YNB+2% glucose, and YNB+2% sorbose media, and the doubling times were calculated by polynomial fits using OD_600_ values after 48, 48, and 144 h incubation, respectively. N=10. Statistical significance was determined by ANOVA and a Dunnett’s post hoc test. **** p<0.0001; *** p<0.001; ** p<0.01; * p<0.05).

Elevated *SOU1* expression could be responsible for survival of the *tlo*Δ/Δ mutants on sorbose medium (72). To determine the relationship between *SOU1* expression and sorbose survival, we constructed heterozygous and homozygous *sou1* deletion mutants in the wildtype background. Surprisingly, the *SOU1/*Δ mutant grew on YNB+2% sorbose, but, consistent with previous reports, the *sou1*Δ/Δ mutant failed to grow under these conditions (Figure S1)(33).

Interestingly, *tlo*Δ/Δ mutants formed colonies on sorbose faster than the *SOU1/*Δ mutant. Colonies appeared on sorbose plates at day 6 for the *tlo*Δ/Δ mutants but not until day 12 for the *SOU1/*Δ mutant. As expected, *SOU1* transcript levels were reduced by approximately half in the *SOU1/*Δ mutant on YPD and were undetectable in the *sou1*Δ/Δ mutant (Figure 2d). However, the expression of *SOU1* varied across a 4-fold range in the *tlo*Δ/Δ mutants when grown on YPD and decreased by approximately half on sorbose-containing medium. Similarly, *SOU1* expression was approximately 50% of the wildtype levels in surviving *SOU1*/Δ heterozygotes on sorbose. This suggests that the variable expression of *SOU1* in *tlo*Δ/Δ mutants on rich medium may contribute to its survival on sorbose by selecting for cells with low *SOU1* expression that is similar to the *SOU1* heterozygote.

Genes besides *SOU1* have been suggested to contribute to sorbose utilization in *C. albicans*. Disruption of sorbose utilization (CSU) genes, such as *PGA56* (formerly as *CSU51*), *CSU52* and *CSU57*, was reported to allow for growth on sorbose-containing medium (54, 73). We constructed clean deletion mutants for each of these three genes individually in both the wildtype and the *tlo*Δ/Δ mutant background to test their influence on sorbose survival. Loss of *PGA56* allowed for growth on YNB+2% sorbose medium; however, neither the *csu52*Δ/Δ nor the *csu57*Δ/Δ mutant grew on sorbose (Figure 2a,b). Mutants for all three *CSU* genes in a *tlo*Δ/Δ background grew on sorbose. These results suggest that the loss of *TLO* alone is sufficient for sorbose survival, and *CSU52* and *CSU57* do not contribute to growth on sorbose. Interestingly, the *pga56*Δ/Δ mutant and the *tlo*Δ/Δ *pga56*Δ/Δ double mutant grew similarly to the wildtype background strain and not the *tlo*Δ/Δ mutant in liquid YPD and YNB+2% glucose media at 30°C (Figure 2c). When grown in liquid YNB+2% sorbose, the doubling time for the *tlo*Δ/Δ *pga56*Δ/Δ double mutant was equivalent to the *pga56*Δ/Δ mutant and significantly faster than the *tlo*Δ/Δ mutant. These results suggest that *TLO* is epistatic to *PGA56* and may promote sorbose survival indirectly through *PGA56*.

To determine if the *TLO* gene family regulates *CSU* genes, we measured their transcript abundance in the *tlo*Δ/Δ mutant, *SOU1* mutants, and the wildtype background strain. Expression of *PGA56*, *CSU52*, and *CSU57* were all significantly reduced in the *tlo*Δ/Δ mutant when grown on YPD at 30°C compared to the wildtype control (Figure 2d). In contrast, transcript abundance of *CSU52* and *CSU57* remained at wildtype levels in the *SOU1*/Δ mutant on YPD medium, although *PGA56* expression was reduced slightly. Complete loss of *SOU1* had no effect on *CSU57* expression on YPD medium, but the expression of *PG56* and *CSU52* decreased 1.5-fold and 4-fold, respectively. When grown on YNB+2% sorbose medium, the expression of all three *CSU* genes decreased further in the *tlo*Δ/Δ mutant v. growth on YPD. Similarly, the expression of all three *CSU* genes decreased in the *SOU1* heterozygote on sorbose medium to levels that were comparable to the *tlo*Δ/Δ mutant. Taken together, this data suggests that *TLO*s activate expression of *CSU* genes on Chr5 and reduced *PGA56* function by either inactivation or modulating gene expression contributes to sorbose survival.

### Tloα and Tloβ directly control *SOU1* expression and sorbose survival

To distinguish between the function of *TLOs* as direct or indirect regulators of sorbose-associated genes, a panel of *TLO* reintegration strains were constructed in the *tlo*Δ/Δ mutants that each contained a single *TLO* gene fused to a C-terminal 6x His-FLAG tag and were integrated at the *NEUT5L* locus (Figure 3a). Introduction of a *TLO*α gene in the *tlo*Δ/Δ mutant restored the wildtype colony morphology; addition of *TLO*β*2* to the *tlo*Δ/Δ mutant induced the formation of smooth colonies with radial filamentation; reintegration of most *TLO*γ genes did not alter the wrinkled colony morphology of the *tlo*Δ/Δ background strain (Figure 3b, S2). The single exception was reintegration of *TLO*γ*4*, which produced smooth colonies identical to strains containing *TLO*α and not *TLO*γ paralogs. When grown on YNB+2% sorbose, strains containing a *TLO*γ gene grew similarly to the *tlo*Δ/Δ parental strain (Figure 3c, S2). However, strains containing a *TLO*α or *TLO*β gene failed to grow in the presence of sorbose, indicating an active role for these *TLO*s in *SOU1* regulation.

**Figure 3.**
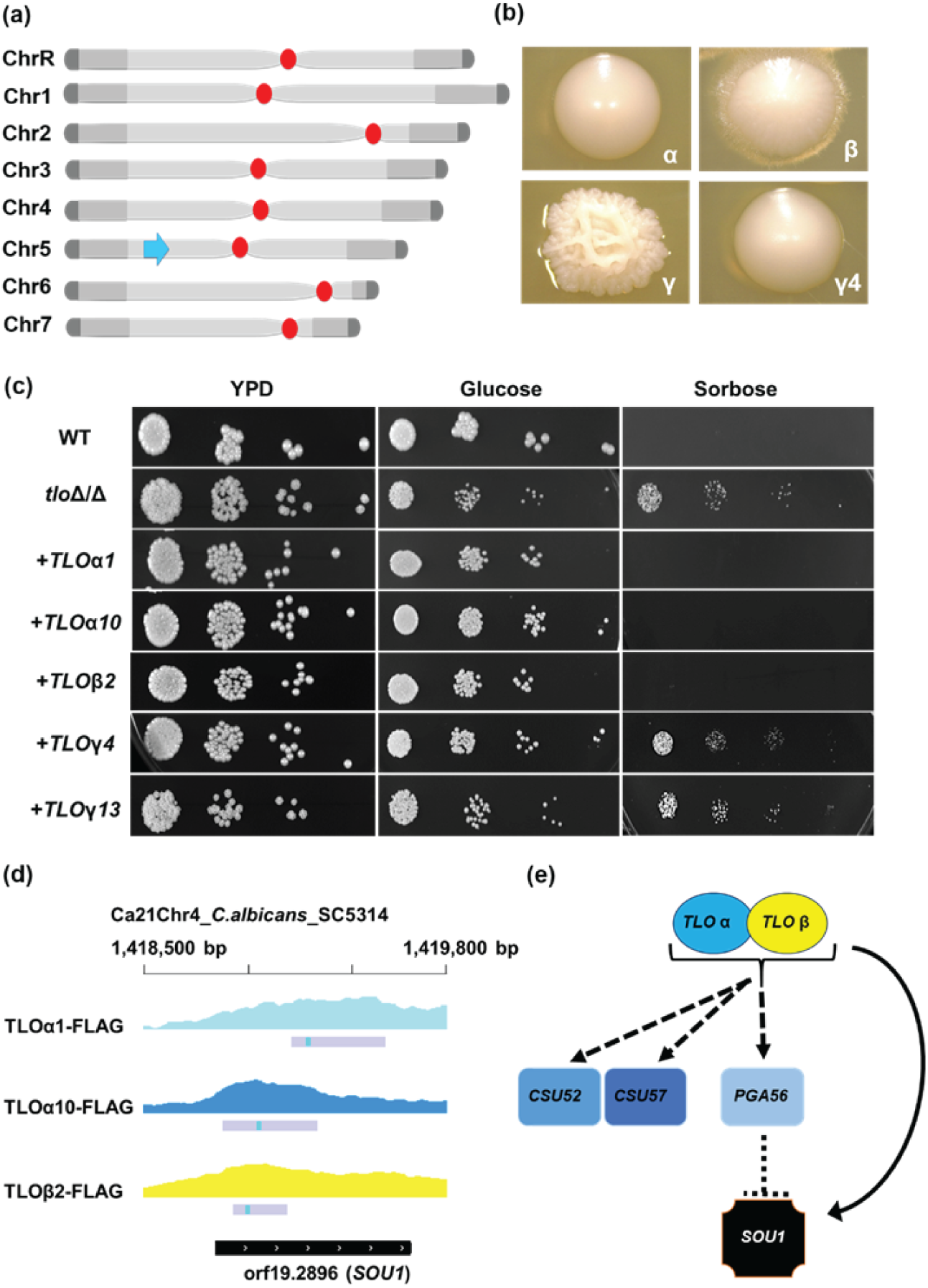
*TLOs* mediate sorbose growth directly and indirectly. (**a**) The *tlo*Δ/Δ mutants were complemented with a single *TLO* gene inserted into the neutral locus *NEUT5L* on Chr5. (**b**) Addition of a single *TLO* gene altered the colony morphology of the *tlo*Δ/Δ mutant that largely fell along architectural groups with the exception of *TLO*γ*4*. (**c**) Cells of the indicated genotypes were spotted in 10-fold dilutions starting with 10^8^ cells on the left onto YPD and yeast nitrogen base (YNB) medium supplemented with 2% glucose or 2% L-sorbose. Colonies were allowed to grow for 2 (YPD) and 6 days (YNB) at 30°C before imaging. (**d**) Tlo-6xHIS/FLAG binding of the *SOU1* locus was determined by chromatin immunoprecipitation and sequencing (ChIP-Seq) normalized for non-specific binding with Nup49-6xHIS/FLAG. Grey bars indicate statistically enriched binding, and the blue box indicates the strongest binding signal. (**e**) Cartoon of *TLO* regulation to produce growth on sorbose via *SOU1* regulation. Dashed lines indicate indirect regulation, and solid lines indicate direct regulation.

We used the epitope tag of reintegrated *TLO* genes to perform chromatin immunoprecipitation (ChIP) of Tloα1, Tloα10, and Tloβ2 binding across the genome. A strain containing a C-terminal 6x His-FLAG tag on the nuclear pore protein *NUP49* served as a control for non-specific binding of nuclear localized proteins. Tagged Tlos varied dramatically in the number of genomic loci bound in each strain. ChIP-Seq identified 683, 182, and 178 regions bound by Tloα1, Tloα10, and Tloβ2, respectively (Table S4). Importantly, all three Tlo proteins bound to the *SOU1* locus (Figure 3d), but no binding was evident at the three *CSU* loci (*CSU52*, *CSU57*, and *PGA56*). ChIP-Seq as also performed with 6x His-FLAG tagged Tloγ4 and Tlo γ13 to determine the specificity of *SOU1* binding. Neither Tloγ bound to the *SOU1* locus, consistent with the lack of sorbose regulation by these genes. Taken together, this suggests that growth on sorbose in the *tlo*Δ/Δ mutant is a consequence of indirect transcriptional activation of the *CSU* genes and direct activation of *SOU1*.

### *TLO*s activate expression of *GAL* utilization genes

To determine the mechanism allowing for growth of the *tlo*Δ/Δ mutants on 2-DOG, we first assessed the stability of the resistance phenotype. As previously noted, cells from the *tlo*Δ/Δ mutant grew on 2-DOG, whereas the wildtype strain lacked any notable growth. Surviving cells from *tlo*Δ/Δ mutant colonies that grew on 2-DOG were allowed to recover on YPD before being replated to both YPD and 2-DOG for fluctuation analysis (Figure 4a). On average, 59% of *tlo*Δ/Δ mutant cells retained viability on 2-DOG, suggesting both a regulatory and genetic mechanism of resistance (Figure 4b).

**Figure 4.**
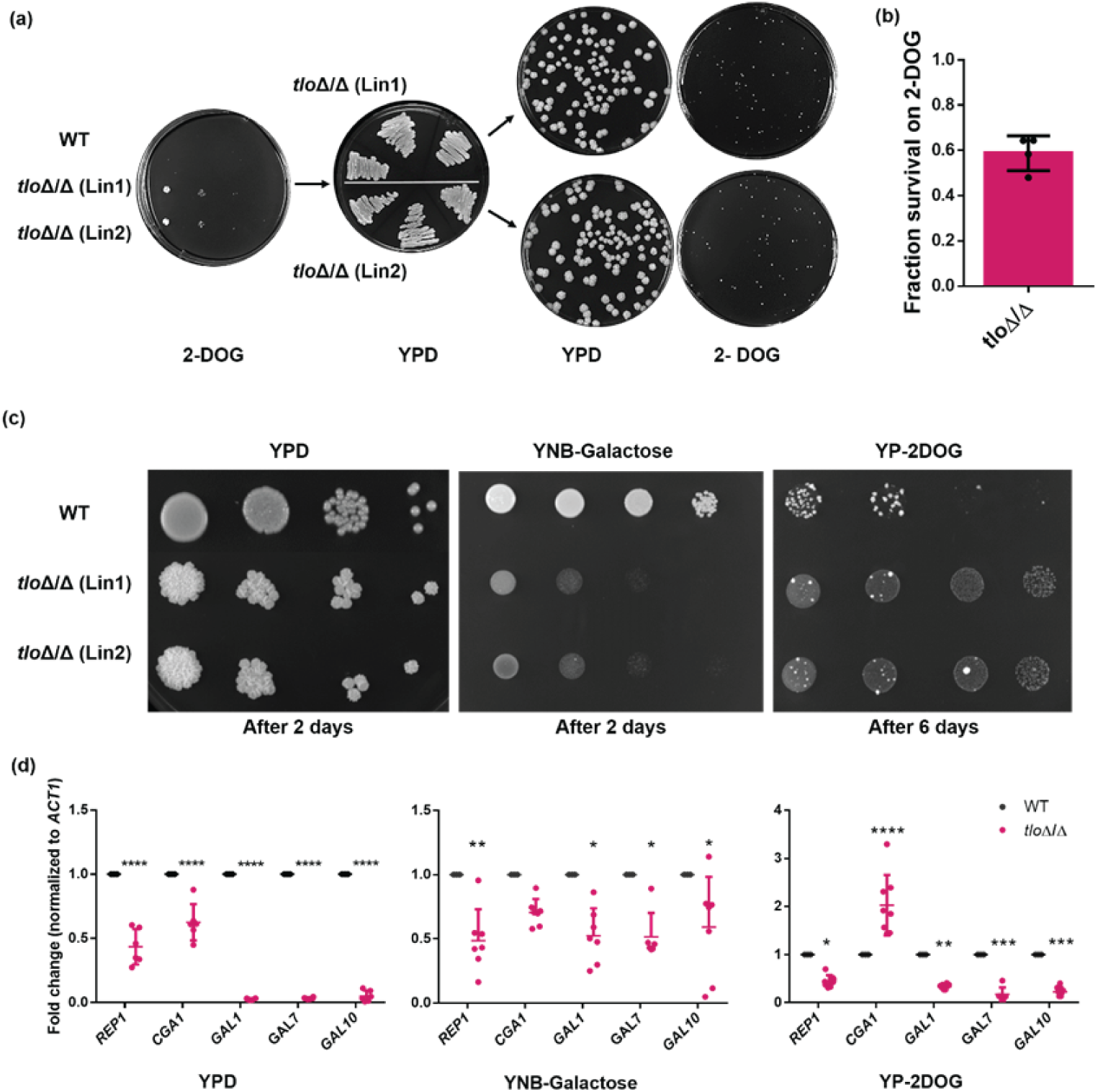
Deletion of *TLOs* disrupts galactose utilization and *GAL* genes expression. **(a**) Fluctuation analysis of cells grown on 2-deoxygalactose (2-DOG) medium. 2-DOG resistant cells were recovered on YPD and subsequently replated to YPD and 2-DOG plates at 100 colonies per plate. Plates were incubated at 30 °C for 5 days. (b) Fraction of *tloΔ*/Δ cells resistant to 2-DOG after replating from YPD medium. (c) 10^8^ cells were spotted in 10-fold dilutions on YPD, YNB with 2% galactose, and YP with 0.2%2-DOG. Colonies were allowed to grow for 2, 2 and 6 days, respectively at 30°C before imaging. (d) Transcript abundance of GAL utilization genes in the wild-type and *tlo*Δ/Δ strains was measured by qRT-PCR after growth as displayed in c). *ACT1* was used for normalization. N=6. Statistical significance was determined by ANOVA and a Dunnett’s post hoc test. **** p<0.0001; *** p<0.001; ** p<0.01; * p<0.05).

To measure the expression of genes associated with galactose metabolism, the growth of the wildtype parental strain and *tlo*Δ/Δ mutants was assessed across media containing either 2% galactose or 0.2% 2-DOG. Interestingly, *tlo*Δ/Δ mutants grew poorly on YNB+galactose but displayed no growth defects on SC or YP media containing galactose (Figure 4c, S3). Wildtype cells did not have a growth defect on YNB+galactose, suggesting that the reduced growth of the *tlo*Δ/Δ reflects a defect in the *GAL* pathway that likely facilitated growth on 2-DOG. Yet, the growth of wildtype and *tlo*Δ/Δ cells on media containing 2-DOG did not reflect the differential growth on media supplemented with galactose as the main carbon source. Both strains grew poorly on YP+2-DOG, neither grew on YNB+2-DOG, and only the *tlo*Δ/Δ mutants grew on SC+2-DOG, the canonical medium used for testing 2-DOG toxicity (Figure S3). Therefore, growth using galactose or 2-DOG was dependent on the base medium supplemented with each carbon source.

Growth of both the wildtype and *tlo*Δ/Δ mutants on YNB+galactose and YP+2-DOG facilitated measuring expression of *GAL*-related genes under identical conditions. The expression of three enzymes in galactose metabolism*, GAL1*, *GAL7*, and *GAL10*, decreased in the *tlo*Δ/Δ mutant regardless of the presence or absence of galactose or the 2-DOG toxic substrate (Figure 4d). Reduced transcript levels of all assayed *GAL* genes in the *tlo*Δ/Δ mutant prompted the interrogation of *REP1* expression. *REP1* ferries the transcriptional activator of galactose metabolism *CGA1* to the promoter of *GAL* genes (74). Transcript levels of *REP1* were also reduced under all conditions in the *tlo*Δ/Δ mutant and decreased to a similar extent as the *GAL* genes in YNB+galactose or YP+2-DOG (Figure 4d). In contrast, *CGA1* did not display consistent patterns of expression like *REP1* or the *GAL* genes. *CGA1* expression decreased on YPD, was unchanged on YNB+galactose, and increased on YP+2-DOG. Taken together, loss of the *TLO* genes decreased *GAL* gene expression, potentially via reduced *REP1* transcript abundance, and this reduced expression likely contributed to the non-heritable source of 2-DOG resistance.

### Loss of *TLO* genes destabilizes the *GAL1* locus

To understand the genetic basis for 2-DOG survival of *tlo*Δ/Δ mutants, we screened colonies for the presence of *GAL1*. Amplification of *GAL1* was successful for most *tlo*Δ/Δ colonies that grew on 2-DOG (Figure 5a). However, three of 25 randomly chosen colonies consistently failed to produce a *GAL1* PCR product when amplifying the coding sequence, suggesting that the gene was absent or mutated to no longer allow for primer binding. The *GAL1* amplicons from twenty 2-DOG^R^ colonies that retained the locus and the wildtype parental strain were submitted for sequencing to assess the allelic composition of the *GAL1* locus. Surprisingly, half (10/20) of the 2-DOG^R^ *tlo*Δ/Δ mutants had undergone loss of heterozygosity at the *GAL1* locus, retaining only the A allele (Figure 5b, S3). The observed loss of heterozygosity could result from replacement of the B allele with a copy of the A allele or completely loss of the B allele. Measurements of *GAL1* copy number by quantitative PCR (qPCR) revealed that many *tlo*Δ/Δ colonies were hemizygous for *GAL1* (Figure 5c). Thus, 2-DOG resistance may be inherited via loss of both *GAL1* alleles, loss of a single *GAL1* allele, or *GAL1* LOH that could be compounded by reduced *GAL1* expression in the *tlo*Δ/Δ mutant (Figure 5d).

**Figure 5.**
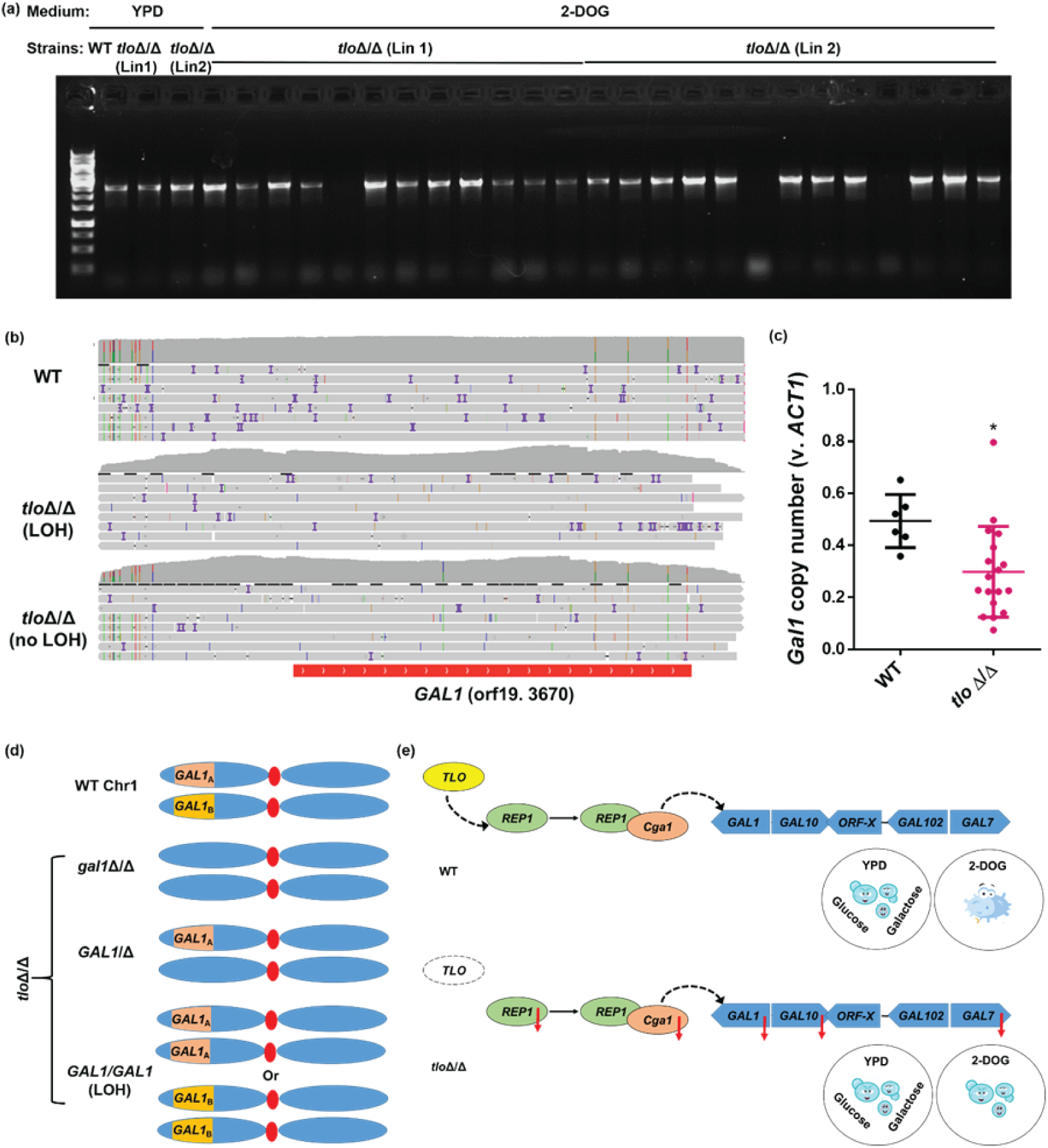
Deletion of *TLOs* promotes *GAL1* copy number variation and loss of heterozygosity. (**a**) PCR amplification of *GAL1* from *tloΔ*/Δ colonies collected from 2-DOG plates. (**b**) Representative read pileups of the wild-type and *tlo*Δ/Δ strains at the *GAL1* locus. Heterozygous positions are shown with colored vertical bars and each color indicates a different base at the position. (**c**) qPCR of *GAL1* from *tlo*Δ/Δ colonies that survived on 2-DOG plates and wild-type control cells on YPD. Copy number was normalized to *ACT1*. N=6(WT),18 (*tlo*Δ/Δ) (**d**) Cartoon of representative *GAL1* outcomes that facilitated survival on 2-DOG medium. (**e**) Model for *TLO* regulation of GAL utilization genes that leads to increased survival on 2-DOG.

### *TLO* mutants display LOH and aneuploidy

Evidence for loss of *GAL1* in specific *TLO* mutants, suggested that the genomes of these strains may be unstable. To assess the influence of *TLO* genes on genome composition, we performed an evolution experiment with the wildtype, *tlo*Δ/Δ mutant, and strains containing a single reintegrated *TLO* gene from each of the three architectural groups. Each strain was plated onto solid YPD medium, allowed to grow at 30°C for two days, and passaged from the thick part of the streak onto new YPD solid medium. After 20 rounds of passaging, 20 individual colonies from each genotype underwent whole genome sequencing (Figure 6a).

**Figure 6.**
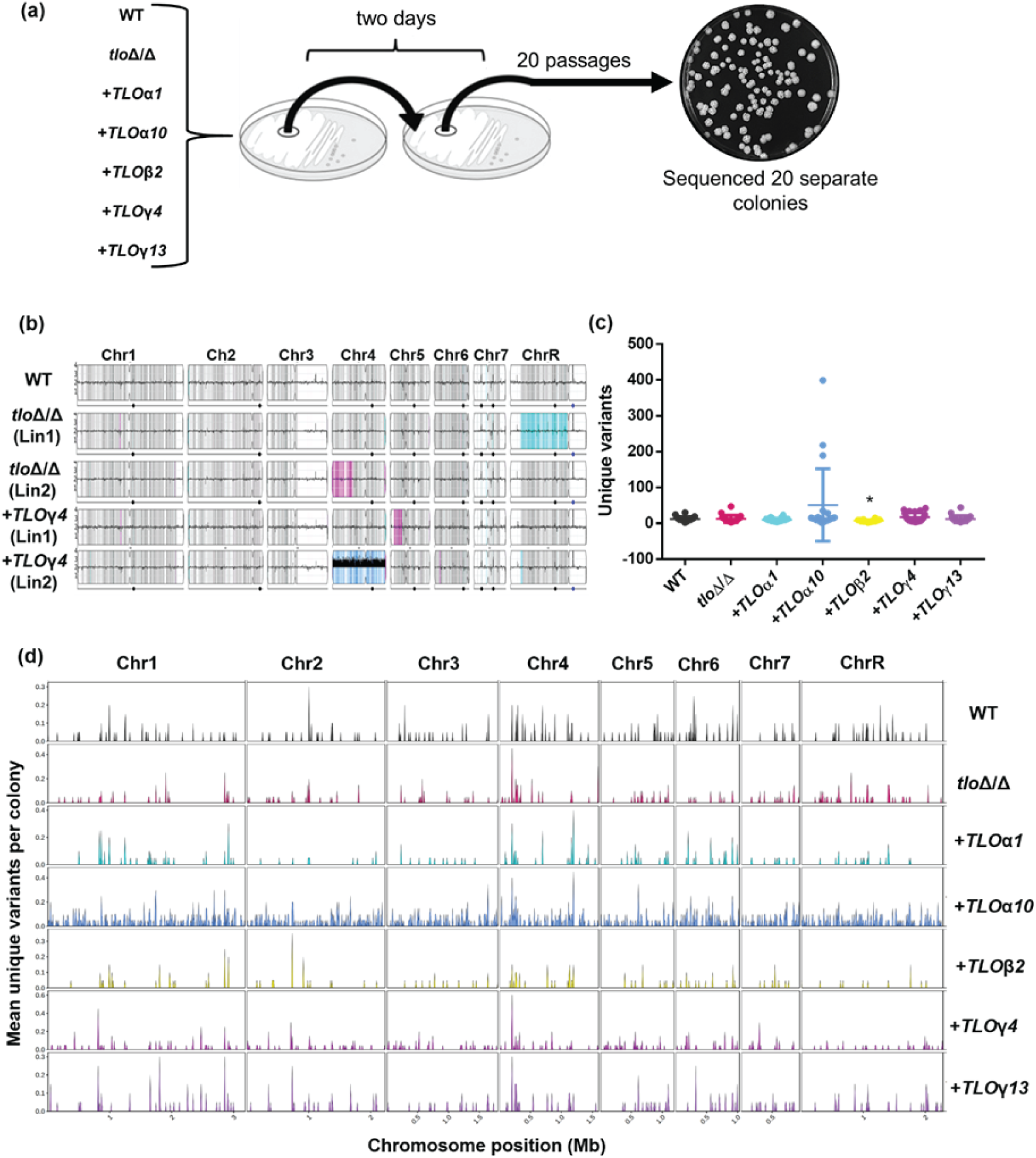
Loss of *TLO*s promotes karyotypic but not genotypic changes. (a) The WT parental strain, *tlo*Δ/Δ mutants, and five *tlo*Δ/Δ strains complemented with different *TLO* genes were passaged every two days on solid rich medium (YPD) agar plates at 30°C. Cells were restruck from the thick part of the streak during each passage. (b) Strain karyotypes were visualized with YMAP. The average ploidy of each chromosome is indicated in black in 10 kilobase bins. Shading indicates genotype where grey=heterozygous, cyan=A alleles, and magenta=B alleles. Blue indicated AAB. (c) Variants in each passaged colony were identified using GATK4, filtered for unique mutations, and plotted by genotype. (d) The unique variants Variants are either shown in aggregate (c) or as classified into bins based on the outcome of each mutational event in each chromosome (f). Statistical significance was determined by Kruskal-Wallis test and Dunn’s multiple comparisons test. (* p<0.05).

Passaging produced some changes in genome composition among the different genotypes. Most strains maintained a heterozygous diploid genome during passaging with the exception of the *tlo*Δ/Δ mutant and *TLO*γ*4*-complemented strain (Figure 6b). Two *tlo*Δ/Δ mutants contained whole or partial chromosome LOH. One *TLO*γ*4* complemented strain had a partial LOH of Chr5, and a second strain was trisomic for Chr4. Approximately 60-80 unique point mutations arose in each passaged strain across the strain genotypes (Figure 6c). A few colonies containing a reintegrated copy of *TLO*α*10* had large increases in the number of new variants following passage, and strains with a reintegrated copy of *TLO*β*2* had significantly fewer new variants compared to the wildtype (average of 44 v. 79, respectively). New mutations existed across all eight chromosomes in each genotype (Figure 6d). The location of new point mutations varied within a single genotype and among genotypes, suggesting that the position of point mutations was not strongly affected by loss of *TLO* genes. Together, this suggests that loss of *TLO* genes could contribute to karyotypic variation that is selected upon during exposure to toxic carbon sources.

### Telomere length is regulated by *TLO* genes via telomere-specific complexes

Whole genome sequencing of passaged *TLO* strains provided an opportunity to define any contributions by *TLO* genes to telomere regulation. Dysregulated telomeres are susceptible to being recognized as DNA double strand breaks and undergoing repeated rounds of breakage-fusion cycles that contribute to chromosome instability (75, 76). Counting the number of reads containing telomere repeats from sequenced wildtype, *tlo*Δ/Δ mutants, and strains with a single reintegrated *TLO* gene provided an estimate of telomere length across the *C. albicans* chromosomes. The genome of *tlo*Δ/Δ mutants had higher frequencies of telomeric repeats compared to the wildtype strain (ANOVA, F(6,113)=21.3, p=1E-4, Figure 7A). Reintroduction of a single *TLO* gene from any of the three architectural groups restored the frequency of reads with telomeric repeats to wildtype levels or reduced them to below the parental wildtype strain when complemented with *TLO*β*2*. To validate these results, telomeric DNA was quantified from the same passaged strains by qPCR. Consistent with the WGS approach, *tlo*Δ/Δ mutants had higher levels of telomeric repeats that generally returned to wildtype levels in the complemented mutants (Figure 7B). Telomeres in the *tlo*Δ/Δ mutant complemented with *TLO*α*1* or *TLO*γ*4* were slightly longer or shorter than the wildtype, respectively.

**Figure 7.**
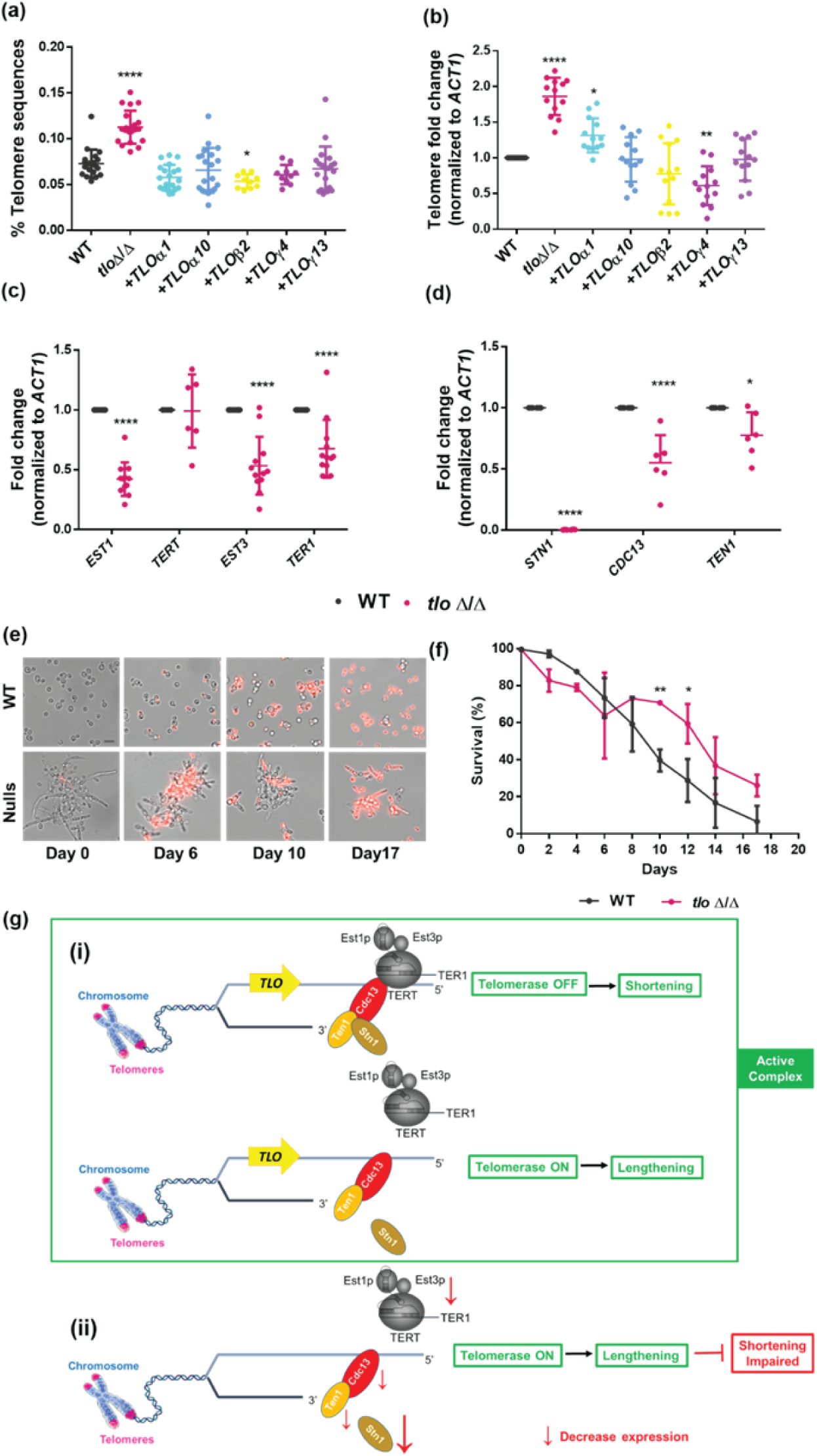
*TLO*s regulate telomere length via Est and the CST capping complex. (**a**) The genomes of 20 isolates of each indicated genotype were sequenced after 21 passages on solid YPD medium at 30°C. The fraction of reads identical to the *C. albicans* telomere repeat is plotted. (**b**) Telomeric repeats were measured by quantitative PCR (qPCR) from DNA of the 20 passaged colonies and normalized to *ACT1*. Transcript abundance of genes in the (**c**) telomerase complex and (**d**) CST capping complex was assessed by qRT-PCR after two days of growth on YPD solid medium at 30°C. The *ACT1* gene was used for normalization. N=8 (**e**) Saturated overnight cultures of wildtype and *tlo*Δ/Δ mutants in liquid synthetic defined complete (SDC) medium were diluted 1:200 into fresh SDC medium and grown at 30°C for the indicated number of days without nutrient replenishment. Cells were stained with propidium iodide (PI) and imaged. Scale bar = 10 µm. (**f**) The percent survival of the wildtype and *tlo*Δ/Δ mutant based on PI staining of 500 cells per replicate. N=4. Statistical significance was determined by one-way ANOVA and a Dunnett’s post hoc test. **** p<0.0001; *** p<0.001; ** p<0.01; * p<0.05). (**g**) Model depicting the mechanism of telomere regulation via CST (Stn1, Ten1, and Cdcl3) regulation of telomerase activity. (i) Wildtype cells control telomere lengths through CST-mediated regulation of telomerase activity to maintain proper telomere lengthening and shortening. In the absence of *TLO*s, (ii) the CST complex members decrease in abundance, leading to increased telomerase activity and longer telomeres.

To determine the underlying cause of increased telomere lengths in the *tlo*Δ/Δ mutant, we investigated the expression of the TERT-associated complex and the CST capping complex that promote and restrict telomere lengthening, respectively (77). The expression of *TERT,* the catalytic telomerase subunit, was unchanged in the *tlo*Δ/Δ mutant (Figure 7C). However, expression of two accessory subunits of telomerase, *EST1* and *EST3*, decreased approximately 2-fold in the *tlo*Δ/Δ mutants along with the telomere template RNA, *TER1*.

An additional complex composed of Stn1-Cdc13-Ten1 (the CST capping complex) is recruited by active telomerase to control telomere length and prevent excessive elongation (78). All three members of the CST complex decreased expression in the *tlo*Δ/Δ mutant (Figure 7D). Importantly, expression of *STN1* was completely ablated in the *tlo*Δ/Δ mutant, which would lead to loss of capping activity and telomere lengthening as observed in the passaged strains.

Increased telomere lengthening can lead to increased cellular lifespan in some contexts (79). *C. albicans* cells grown under nutrient starvation conditions will ultimately undergo cellular senescence and death. We hypothesized that *tlo*Δ/Δ mutants would have extended viability under starvation conditions as a result of increased telomere lengths. To test this hypothesis, wildtype and *tlo*Δ/Δ mutant cells were grown in liquid SCD at 30°C for 17 days without nutrient replenishment. Nearly all cells in the wildtype and the *tlo*Δ/Δ mutant were viable at day 0 (Figure 7e,f). However, both strains displayed an increasing number of dead cells, as detected by staining with propidium iodide, over time. By day 8, the *tlo*Δ/Δ mutant began to display improved viability compared to the wildtype that extended through the end of the experiment (Figure 7f). Thus, *TLO* genes control telomere length through telomerase activity and promote cell viability over extended periods of starvation, consistent with having longer telomeres.

## Discussion

Survival of *C. albicans* in the mammalian host is dependent on their ability to respond and adapt to the surrounding environment. These responses can be mediated by transcriptional and genome plasticity to produce changes that facilitate survival and be passed on to future generations. The phenotypic consequences of *TLO* loss reveal an unexpected connection between transcriptional regulation and genome stability in *C. albicans*. The initial observation of *tlo*Δ/Δ mutant survival on toxic carbon sources suggested that transcriptional regulation of sorbose and galactose metabolism promoted growth, but a deeper analysis indicated that *TLO* loss is accompanied by increased genome instability that also contributes to survival. *TLO* genes exert pleiotropic effects on maintaining genome integrity by regulating recombination processes associated with LOH and telomere homeostasis as subunits of the Mediator complex. Yet, *TLO*-dependent genome regulation differs among individual paralogs and generally falls along architectural group boundaries. Most phenotypes were restored by reintroduction of α-and β-clade *TLO*s but not *TLO*γ genes, with the exception of telomere length maintenance. Together, these findings demonstrate that *TLO*s contribute to phenotypic variation by direct regulating gene expression and constraining genome instability.

The genetic basis for genome integrity is poorly described. Most frequently, genes that promote these phenotypes are associated with the cell cycle, DNA segregation, and cell division. The involvement of Mediator in regulating these processes is less surprising when considering its role in global transcriptional regulation. Mediator has previously been implicated in supporting DNA repair processes by either activating gene expression of repair proteins or direct regulation of repair proteins (80–83). Our experiments suggest Mediator limits CIN in *C. albicans* by suppressing recombination and telomere dysfunction. This could occur alongside of genes involved in DNA repair or, in the case of recombination, be a direct result of this regulation. Restricting telomere activity would prevent a positive feedback loop of genome instability that leads to genome fragmentation and cell death (84, 85).The initial drop in cell survival among the *tlo*Δ/Δ strains during starvation may reflect an increased CIN and cell death phenotype that is compensated for through longer telomeres that improve survival during starvation.

The most common mechanism of survival on sorbose medium involves chromosome 5 monosomy, which reduces the dosage of negative regulators of *SOU1* expression (35, 72). Our results are consistent with a previous report that loss of *PGA56* on Chr5 can directly improve sorbose survival (35). In contrast to prior work, loss of *CSU57* or *CSU51* did not provide resistance to sorbose (35, 73). The basis for this difference is not immediately clear although the previously constructed *csu57*Δ/Δ mutant was built in a *csu51*Δ/*CSU51* background that could affect the phenotype. Furthermore, the original identification of *CSU51* as a sorbose-utilizing gene was completed prior to construction of the *C. albicans* genome and could be confounded by undetected chromosomal changes that are known to be present in strain CAF2-4 (86). Use of the *tlo*Δ/Δ mutants clarifies the pathway that leads to sorbose resistance, which includes *PGA56* and the potential direct control of *SOU1* expression by Tlo. It is tempting to speculate that ergosterol depletion observed in strains monosomic for Chr5 may similarly be a consequence of reduced expression of Tlo regulatory targets that could be shared between these phenotypes since both sorbose and ergosterol phenotypes exhibit the same *TLO*α/β-clade dependence (21, 55).

Transcriptional repression or loss of *GAL1* allowed for survival on 2-DOG in our *tlo*Δ/Δ mutants. At the transcriptional level, reduced expression of *REP1* in the *tlo*Δ/Δ mutants would limit the availability of this regulatory scaffold to recruit the Cga1 activator to *GAL* promoters in the presence of galactose, leading to reduced *GAL* gene expression (Figure 5e) (74). Loss of *GAL1* that is observed in roughly half of the 2-DOG resistant colonies would further reduce transcript levels and facilitate survival. Complete loss of *GAL1* was less frequent than loss of a single copy. Localized recombination events could remove a single copy by unequal exchange between chromosome homologs. A subsequent LOH event could produce *gal1*Δ/Δ strains with 2-DOG resistance. Regardless, the ability to grow on 2-DOG is likely the outcome of multiple simultaneous regulatory and genetic changes following loss of the *TLO* genes.

Telomere length reflects a balance between multiple interacting forces to maintain chromosome integrity. This balance is upset in the *tlo*Δ/Δ mutants, resulting in longer telomeres likely resulting from altered telomerase-mediated elongation and CST-complex capping activity that restrains elongation (87). Reduced expression of the accessory telomerase subunits *EST1* and *EST3* will upset telomere length control (88), which will be exacerbated by reduced expression of CST complex members. However, we cannot rule out a direct role for telomere binding by Mediator that is mediated through the Tlo subunit as observed for the Tail module in yeast (89). Improved viability under starvation conditions accompanied the elongated telomeres in the *tlo*Δ/Δ cells is broadly consistent with links between telomere maintenance and chronological lifespan in yeast (65).This does not imply that increased telomere length always lead to improved survival as forced telomerase tethering elongates telomeres in *S. cerevisiae* but increased genome instability and shortened chronological lifespan (79). Interestingly, the restoration of telomere length by representative α, β, and γ *TLO* architectural groups identified telomere homeostasis as a broadly *TLO*-dependent phenotype. This contrasts sharply with the clade-dependent phenotypes for sorbose and 2-DOG survival. We anticipate this is not a consequence of their subtelomeric position on the *C. albicans* chromosomes since each *TLO* was reintegrated at the same chromosome-internal position.

A growing body of evidence suggests that *TLO* function is largely separated by architectural groups. As observed in recent work, *tlo*Δ/Δ mutant phenotypes here are generally complemented by the addition of a *TLO*α or *TLO*β gene but not the Tloγ paralogs (19, 21). This may suggest that *TLO*γ genes lack function in *C. albicans* but is refuted by two pieces of data in the present study. First, telomere homeostasis could be restored by addition of any *TLO* gene in the *tlo*Δ/Δ background. Restored function could reflect a conserved *TLO* function that is independent of Mediator or evidence for Tloγ incorporation into Mediator and divergence in their downstream regulons. While Tloγ members have not been identified in association with Mediator (79, 90), their low expression compared to other *TLO* architectural groups may obscure their presence in the complex (15, 17). Alternatively, Mediator-independent functions in gene regulation have been observed for the Tail module in *S. cerevisiae*, which includes the *MED2* homolog to *C. albicans TLO* genes (91). The second piece of evidence for *TLO*γ function is the restored 2-DOG sensitivity of the strain containing *TLO*γ*4* that phenocopied the addition of *TLO*α/β genes. Yet, *TLO*γ*4* is truncated more extensively than other *TLO*γ genes by a second LTR insertion and could have a unique functional profile because of its altered structure compared to members of the other three architectural groups.

*C*ontributions to *C. albicans* adaptation through transcriptional regulation and genome stability by *TLO* genes suggests a larger role for Mediator in phenotypic responses. In addition to the well documented role of Mediator subunits in transcriptional control of target loci, these subunits can regulate processes that control genome architecture to throttle the generation of new genomic and phenotypic variants. In *C. albicans*, the broad impact of this regulation is observed on gene loss or regulated expression and altered telomere homeostasis. Together, these findings identify *TLO*s as important regulators of both transcriptional plasticity and genomic stability that can provide two simultaneous mechanisms for *C. albicans* can adapt to changing environmental conditions.

## ACKNOWLEDGEMENTS

We would like to thank members of the Anderson lab for fruitful discussions in designing this work and its format for presentation. Generative AI was not used in any part of this study from design to interpretation to construction of this manuscript.

## COMPETING INTERESTS

These authors declare no competing interests.

## AUTHOR CONTRIBUTIONS

Conceptualization: ZS and MZA. Data curation: ZS, EM, and CCC. Formal analysis: ZS, PSH, EM, and CCC. Funding acquisition: MZA. Investigation: ZS, PSH, ALW, AA. Methodology: ZS, PSH, EM, CCC, ALW, and MZA. Project administration: MZA. Resources: MZA. Supervision: MZA. Validation: ZS. Visualization: ZS, CCC, and MZA. Writing: ZS and MZA.

## FUNDING

This work was supported by NSF CAREER Award 2046863 and the Chan Zuckerberg Science Diversity Leadership Award to MZA. This work was also supported by National Institutes of Health Award 1F31AI167576-01A1 to ALW.

## Notes

### Competing Interest Statement

The authors have declared no competing interest.

