## Supplemental Materials for "A Mediator subunit regulates genome stability via direct and indirect transcriptional targets in *Candida albicans*"

### Supplement

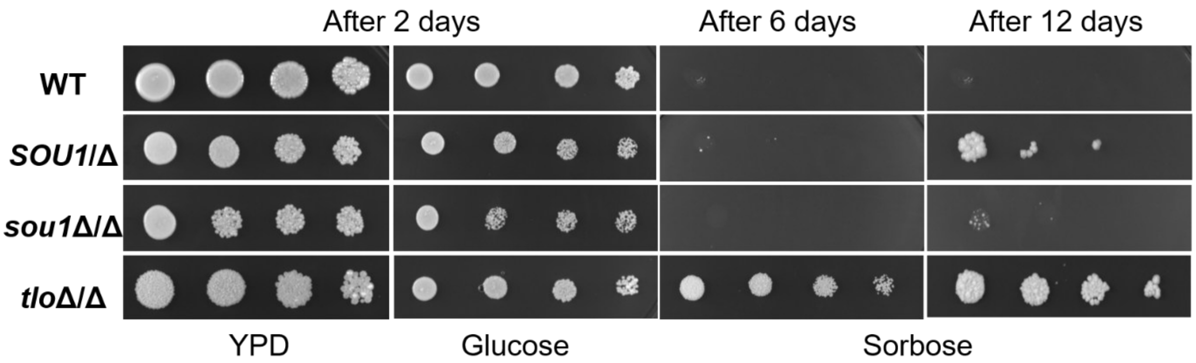


**Figure S1. Differential growth of the *tlo*Δ/Δ and *SOU1*/Δ mutant on sorbose.** Cells were spotted in 10-fold dilutions starting with $10^8$ cells on the indicated media in 6 µL. Colonies were allowed to grow for 2, 6, or 12 days at 30°C before imaging.

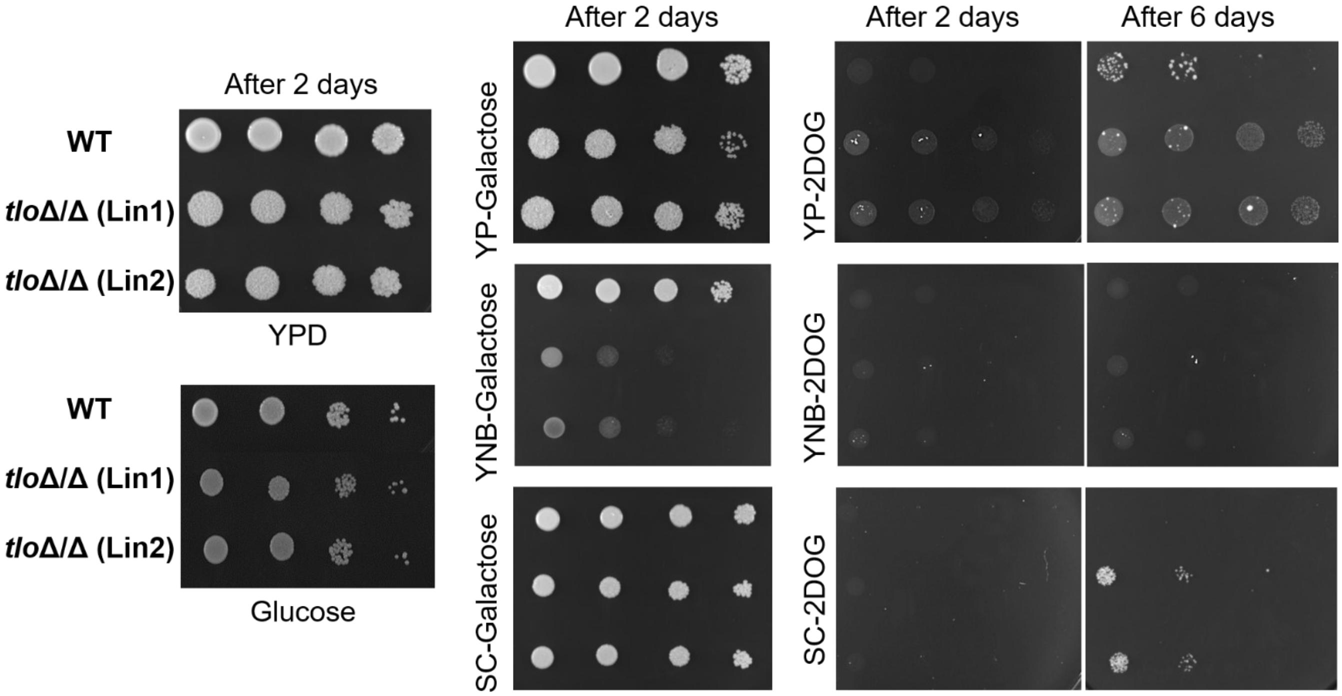


**Figure S2. Nutrient context and *TLO* genotype determine galactose utilization phenotypes**. Cells were spotted in 10-fold dilutions starting with $10^8$ cells on the indicated media. Colonies were allowed to grow for 2 or 6 days at 30°C before imaging as indicated.

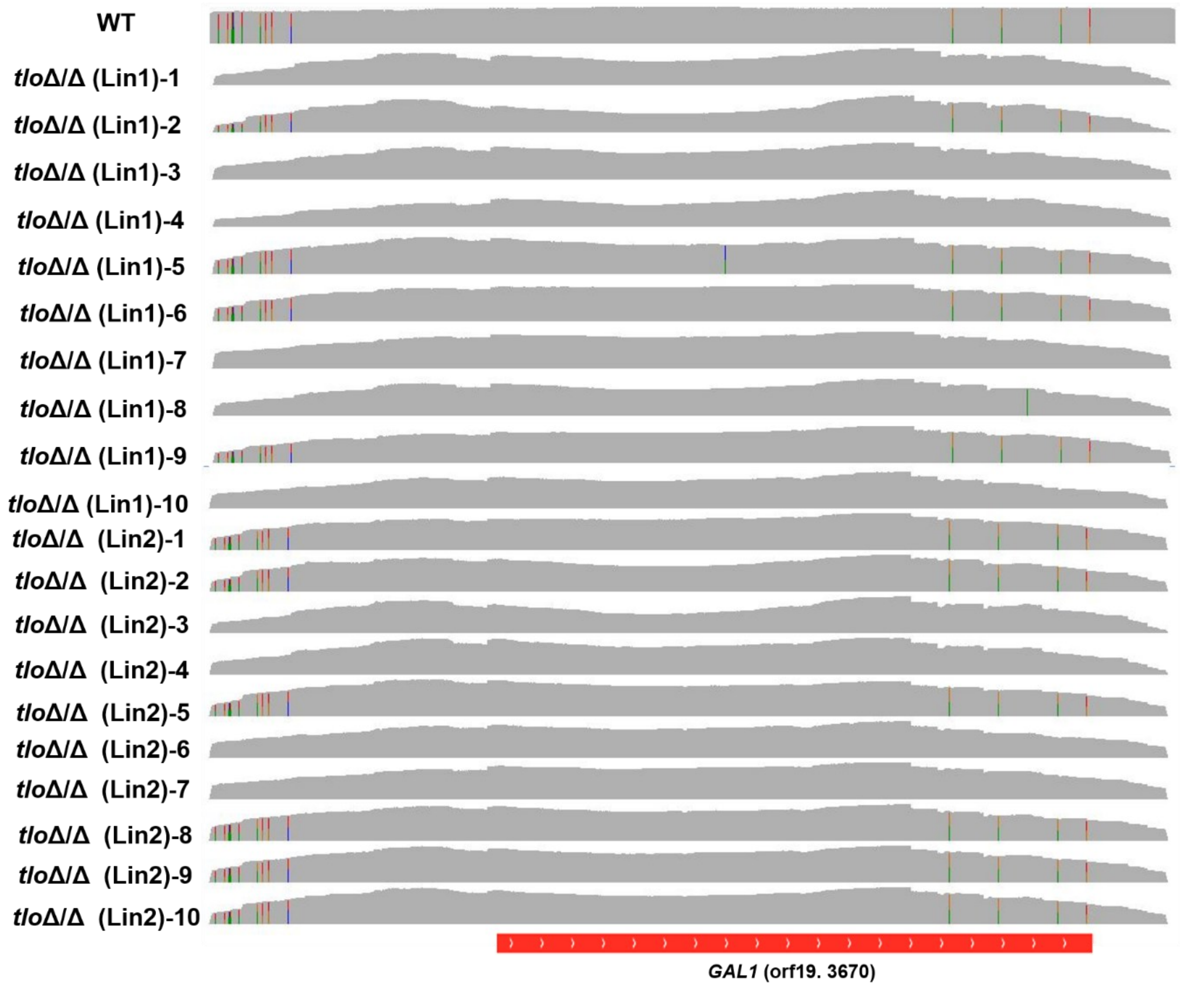


**Figure S3. Biased loss of the *GAL1* A allele in 2-DOG resistant *tloΔ/Δ* mutants.** The *gal1* locus of twenty colonies from 2-DOG plates was PCR amplified and sequenced. The data was visualized using IGV. Heterozygous positions are shown with colored vertical bars and each color indicates a different base.

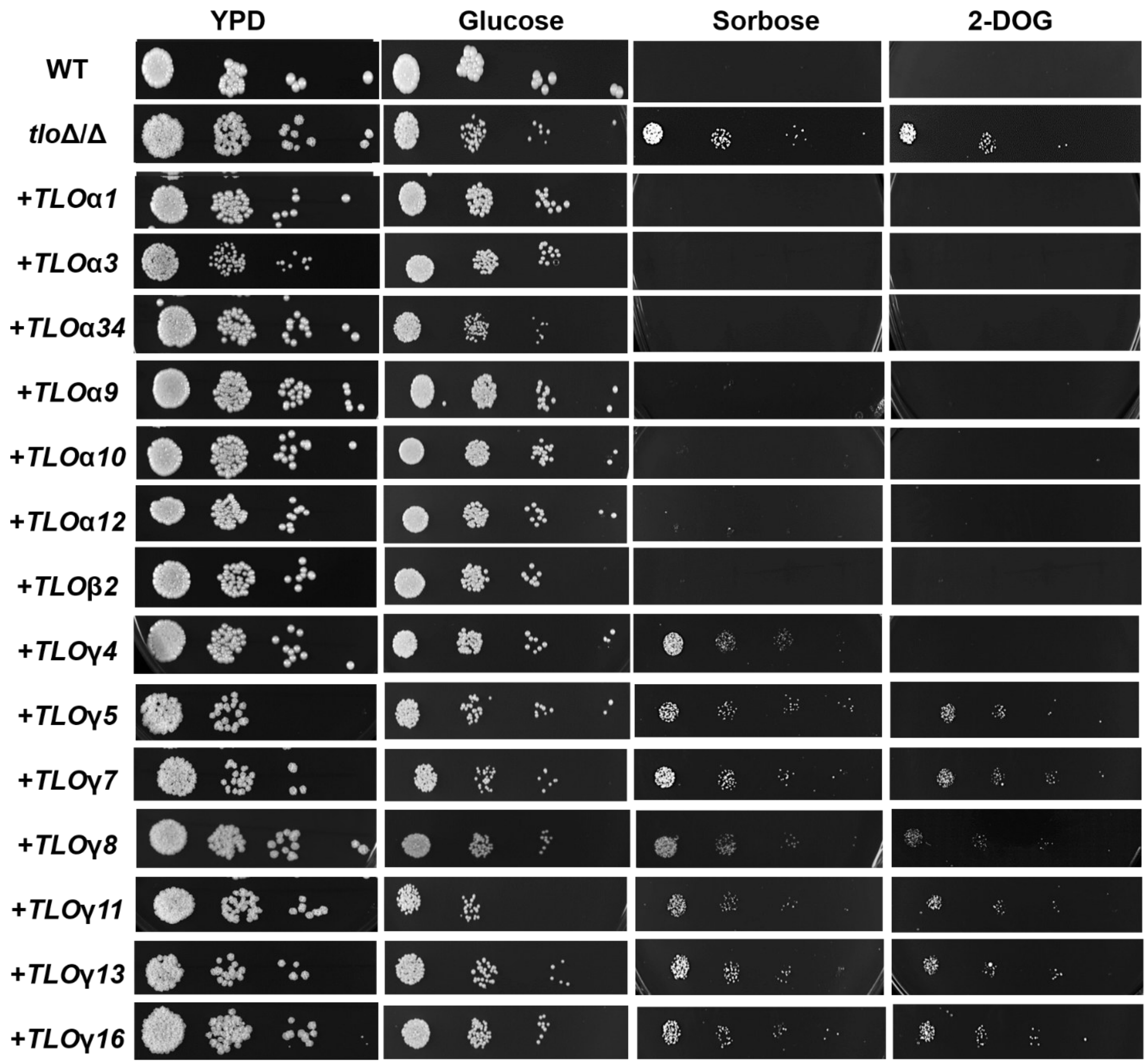


**Figure S4. Functional complementation with *TLOs* produces different growth phenotype.** Cells were spotted in 10-fold dilutions starting with $10^6$ cells onto YPD and yeast nitrogen-based medium supplemented with glucose, 2% L-sorbose, or synthetic complete medium (SCD) supplemented with 0.2% 2-DOG. Colonies were allowed to grow for 5 days at 30°C before imaging.

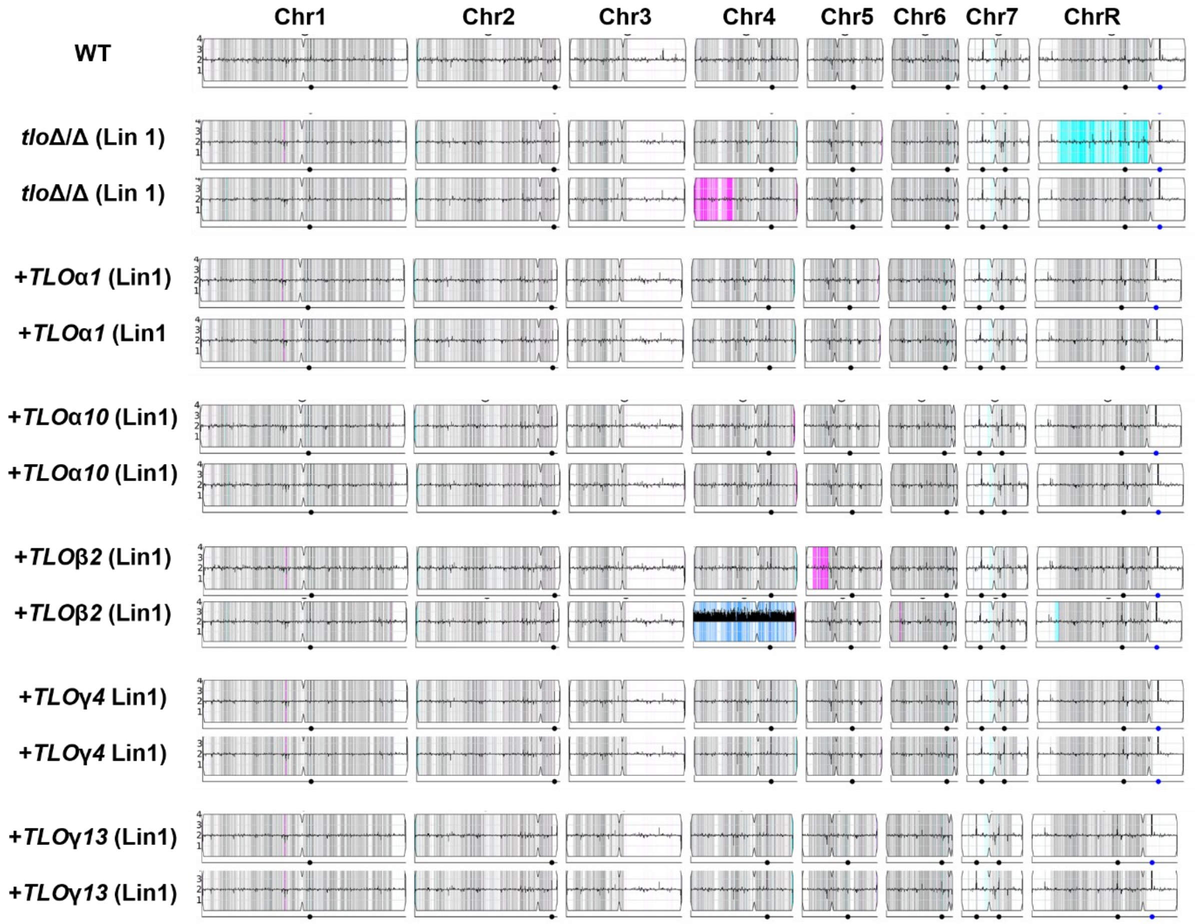


**Figure S5. Karyotypic variation in passaged *TLO* mutants.** Twenty independent lines from the WT and 10 independent lines of each *tlo*Δ/Δ mutant strain and their corresponding reintegrant strains were passaged every two days as populations on YPD medium at 30°C. After 21 passages, DNA was isolated and sequenced and karyotypic changes visualized in YMAP (Abbey et al., 2014). The height of the solid black bars indicates copy number in 10 kilobase bins (flat black line = 2N). Grey, cyan, and magenta colors represent heterozygous, homozygous homolog A, and homozygous homolog B regions respectively. Blue indicates A/A/B allelic balance, and purple indicates A/B/B allelic balance. The centromeres are indicated by pinching in the plot. The ribosomal DNA locus and the Major Repeat Sequence (MRS) positions are indicated by blue and black dots below each plot, respectively.
